# Rapid and specific detection of single nanoparticles and viruses in microfluidic laminar flow via confocal fluorescence microscopy

**DOI:** 10.1101/2023.12.31.573251

**Authors:** Paz Drori, Odelia Mouhadeb, Gabriel G. Moya Muñoz, Yair Razvag, Ron Alcalay, Philipp Klocke, Thorben Cordes, Eran Zahavy, Eitan Lerner

## Abstract

Mainstream virus detection relies on the specific amplification of nucleic acids via polymerase chain reaction, a process that is slow and requires extensive laboratory expertise and equipment. Other modalities, such as antigen-based tests, allow much faster virus detection but have reduced sensitivity. In this study, we report the development of a flow virometer for the specific and rapid detection of single nanoparticles based on confocal microscopy. The combination of laminar flow and multiple dyes enable the detection of correlated fluorescence signals, providing information on nanoparticle volumes and specific chemical composition properties, such as viral envelope proteins. We evaluated and validated the assay using fluorescent beads and viruses, including SARS-CoV-2. Additionally, we demonstrate how hydrodynamic focusing enhances the assay sensitivity for detecting clinically-relevant virus loads. Based on our results, we envision the use of this technology for clinically relevant bio-nanoparticles, supported by the implementation of the assay in a portable and user-friendly setup.

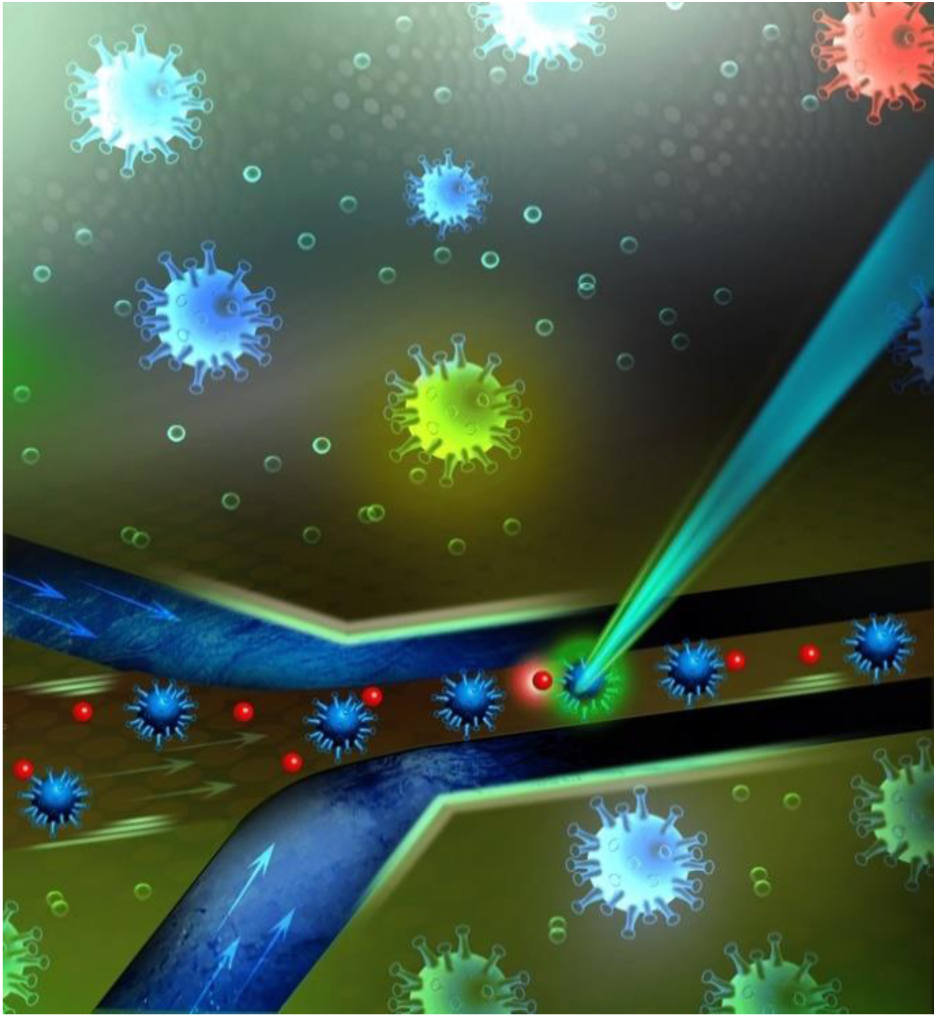

## Introduction

The sensitive and rapid detection of small-scale particle such as bio-nanoparticles (bio-NPs), viruses, liposomes, extracellular vesicles of specific biomolecules plays a crucial role in diagnostics and treatment. The relevance of such methods has become clear during the SARS-CoV-2 pandemic^1^. Mainstream virus detection relies on (i) antigen detection^2–4^, (ii) nucleic acid amplification via polymerase chain reaction (PCR)^5–7^ or (iii) serological tests, which can detect antibodies post infection^8,9^. Rapid antigen-based methods have been used extensively for the detection of SARS-CoV-2. Yet, they feature low accuracy and high false negative rates, and hence serve primarily as an initial screening step^4^. In PCR-based detection, which is highly sensitive and accurate, the viral infection can be verified indirectly via enzyme-based amplification of genetic material. However, this approach is laborious with infection verification results reported hours after initiation of testing^5^. Furthermore, non-nucleic acid biomarkers cannot be amplified via PCR-based methods, limiting its scope.

Other established methods rely on flow-cytometry (FCM) equipment to directly detect virus particles via light scattering and/or fluorescence signals and were also dubbed flow “virometry”^10^. Since virus particles are small and optically transparent, light scattering signals are weak and hence difficult to observe with common FCM equipment. By combining different fluorescent dyes and immunolabeled fluorescent antibodies for labeling different molecular properties of viruses, (e.g., nucleocapsid, matrix, glycoprotein, viral envelope and genome), one can analyze viruses by FCM^10–12^. However, it is necessary to apply data gating to exclude larger cells and non-specific fluorescence signals reducing the sensitivity of this approach for unstained or weakly-stained viruses^10,13^. Additionally, FCM requires high particle loads for distinguishing the correct subpopulation, which might obscure clear identification of distinct viruses. An attractive approach for attaining single virus counting is to use confocal fluorescence microscopy, similar to detection of freely-diffusing particles in fluorescence correlation spectroscopy (FCS), for direct virus detection in microfluidic flow channels. Recent attempts to achieve this have been reported by Niu, Ma *et al.*^14^. In this attempt, both light scattering from virus particles and fluorescence from nucleic acid staining of single particle signal bursts were recorded in a 25 femtoliter (fL) probe volume. Since there are many bio-NPs that contain nucleic acids (e.g., different viruses, exosomes) it is unclear whether nucleic acid staining provides the best means to detect specific virus particles in a heterogenous sample. Additionally, the probe volume that is used, which is the confocal volume, can be as low as 1 fL, which can enhance the sensitivity via improved signal-to-noise ratio. Confocal-based single virus particle detection in minutes by counting coincident bursts was also achieved in assays targeting nucleic acids non-specifically^15^ and later using machine-learning approaches to achieve the basis for the identification of specific viruses^16^.

In this work, we developed a confocal-based single virus particle detection assays, with which we identify specific viruses in minutes by counting coincident bursts with high specificity based on specific antigen interactions and particle volume.

Based on the existing limitations and considerations, we here describe the development of a sensitive small-scale particle counting assay for the optical detection of viruses. For an unequivocal virus identification we observe acoincident signals in microfluidics-based laminar flow. The result combine the ability to identify particle volumes and surface protein epitopes. We demonstrate the ability of the approach to count a sufficient number of single particles with diameters ≥100 nm, one at a time, within minutes. In addition, we demonstrate the ability of the herein assay to specifically detect virus particles SARS-CoV-2 and the rVSV-ΔG-spike, a recombinant vesicular stomatitis virus (rVSV) genetically engineered to express the SARS-CoV-2 spike protein^17^ within minutes. We finally show that hydrodynamic focusing can assist in improving the particle counting sensitivity to a concentration regime of ∼10^4^ particles/mL, which corresponds to realistic virus loads in bodily fluids. The assays presented in this work were performed using both a lab-based confocal microscope and a portable minimalistic 3D-printed microscopy confocal setup^18^. Therefore, the herein described flow virometer can be envisioned as deployable for rapid screening for viruses

## Results

### Direct nanoparticle detection

The assay presented here detects single small-scale particles (>100 nm) in microfluidic laminar flow based on the correlated observation of fluorescent signals related to the virus, its volume, and specific antibody labels on its capsid^19^. Our assay was inspired by inverse fluorescence cross-correlation spectroscopy (iFCCS)^20,21^, in which high dye concentrations are used for counting cross-correlated events of signals from single small-scale particles. In iFCCS, freely-diffusing dye molecules, such as fluorescein, fill the probe volume of a confocal microscope to yield a constant fluorescence signal, which we refer to as the *nonspecific signal*. Large particles (e.g., polystyrene beads, viruses) give rise to abrupt changes to the nonspecific signal, i.e., dips or bursts. These serve as indicators for the presence of small-scale particles in the sample.

As a particle traverses the probe volume, the nonspecific signal decreases due to the exclusion of free dye molecules by the particle volume (Fig. 1A). For such a particle to be detected, the change of the nonspecific signal should be sufficiently large relative to the amplitude of the noise around the mean nonspecific signal. To reduce the heterogeneity of the sizes and durations of the changes in the nonspecific signal, we performed all experiments under laminar flow conditions within a commercially-available microfluidic channel (Figs. 1D, S1A,B). Minimizing the variations in both the durations and sizes of changes in the nonspecific signal is crucial for strengthening the direct relation between signal changes and particle volumes. This contributes to robust identification of signal changes, and to obtain higher particle count rates (Figs. S1C, S2).

**Fig. 1.**
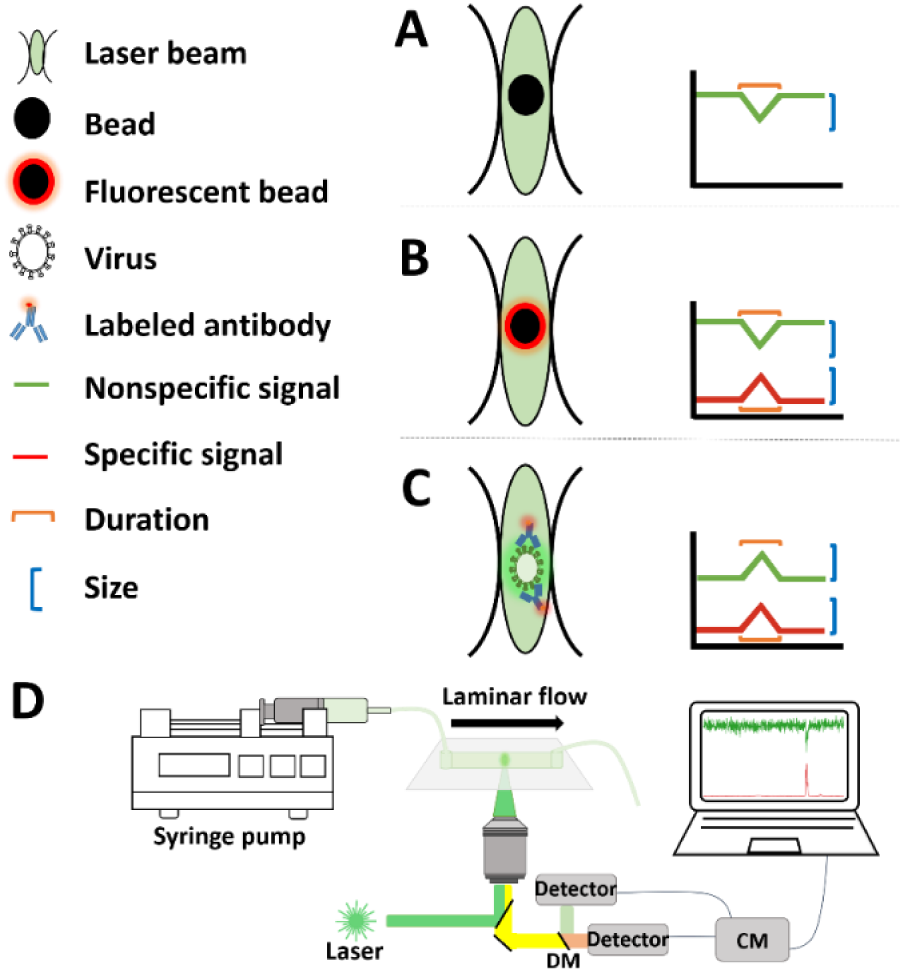
Nanoparticle identification concept based on volume and specific binding. Determining nanoparticle (NP) volume as a function of duration and size of dips (A, B) or bursts (C) in the nonspecific signal (i.e., the fluorescence signal from the free dyes in solution) as the NP traverses the probe volume, i.e. the confocal volume at the focus of the laser beam. **A-C**. Finding the volume of unlabeled beads (black spheres; **A**), the volume of a of a specific NP using red-labeled beads (red-black spheres; **B**), or specific virus particles using antibodies (red fluorescent labeled; **C**), in the presence of a high concentration of free dye. **D**. Schematics of the experimental setup, where excitation light (green) is focused by an objective lens into the analyte flowing in a laminar flow (horizontal arrows) within a microfluidic channel mounted on a glass coverslip. Constant flow rate is achieved using a syringe pump. Fluorescence from free fluorescein dyes (nonspecific signal) and red-labeled dye (specific signal) is collected through the same objective lens, then spectrally split, detected using two detectors and observed as signal changes in the acquisition computer (coincident dip in green nonspecific signal and burst in red specific signal). CM - counting module; DM – dichroic mirror.

We first tested the concept of the detection of fluorescence reduction in the nonspecific signal using unlabeled polystyrene beads with different diameters in the presence of 500 µM fluorescein at different flow rates (Fig. S2) within a microfluidic channel with a 100x1,000 µm^2^ cross-section area, using a confocal microscope (details see *Methods*). When polystyrene beads traverse the probe volume, a temporal reduction in the constant nonspecific signal occurs, forming a signal *dip* (Figs. 1A, 2A). The sizes and durations of the signal dips per bead diameter were recorded, and their distributions were calculated (Figs. 2, S2). Differences between the histograms of sizes and durations of the observed dips validated the ability of the assay to distinguish between small-scale particles of different diameters (Fig. 2). In fast flow rates the variance of dip sizes and durations decreased, however, the distinguishability between the different particles diameter reduced as well (Fig. S2). Accordingly, we selected a flow rate of 750 nL/min (2.08x10^-6^ m/s calculated mean velocity in the microfluidic channel, see Eq. S3) for further experiments, providing optimal discrimination between particles of different diameters.

**Fig. 2.**
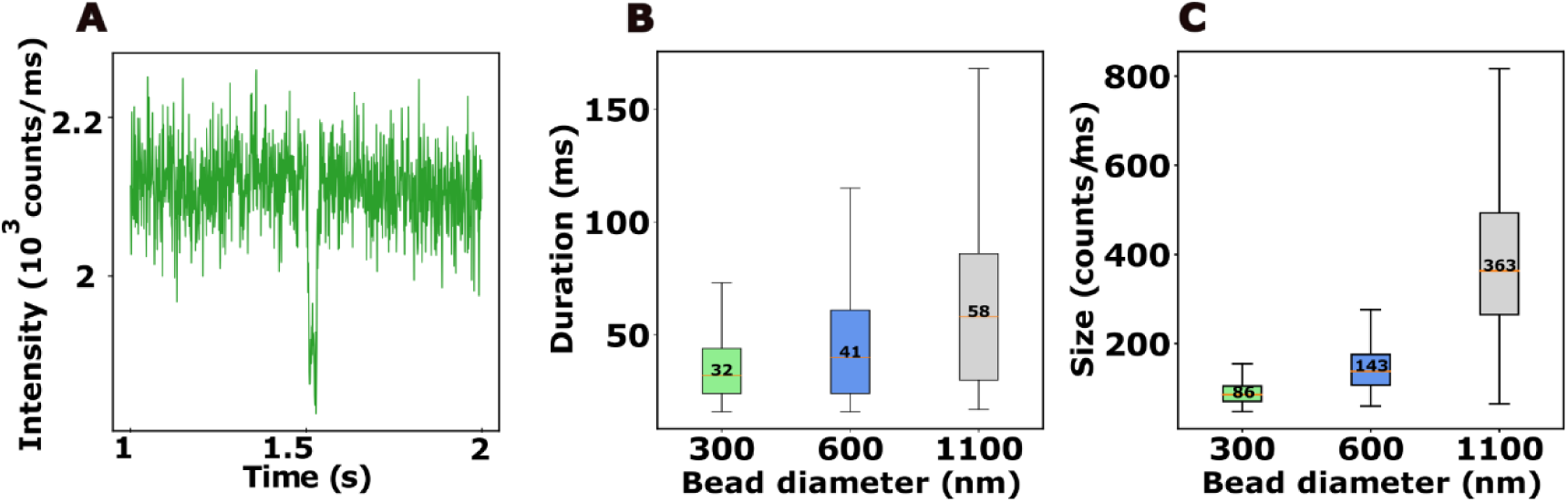
Signal properties versus bead diameters. **A**. A dip in the nonspecific signal caused by the flow of spherical beads with diameters of 600 nm through the probe volume in a 1 s time window (similar examples in Fig. S3). **B, C.** Distributions of nonspecific signal dip sizes and durations from 7 min. acquisitions of beads with diameters of 300, 600 and 1,100 nm (accumulation of 246, 881 and 470 detection events from 4, 6 and 4 different repetitions, respectively, see Fig. S2) with a constant flow rate of 750 nL/min (2.08x10^-6^ m/s calculated mean velocity in microfluidic channel, see Eq. S1). P-values=*** between samples of different beads size, see Table S1. The orange horizontal line represents the median value.

Using the optical setup and assay described above, we detected beads with diameters ≥300 nm. To facilitate the detection of smaller particles as well as to improve the applicability and affordability of the detection method, we employed a recently introduced adaptable microscopy platform dubbed Brick-MIC^18^ (Fig. 3A). Importantly, using PMTs in the 3D-printed setup instead of the more sensitive hybrid PMTs (as in the setup in Fig. 2), allows recording larger signals due to higher detection saturation levels. This, in turn, allow reducing signal variances for larger signals and therefore enhances the signal-to-background ratio in our signal change detection procedure.

**Fig. 3.**
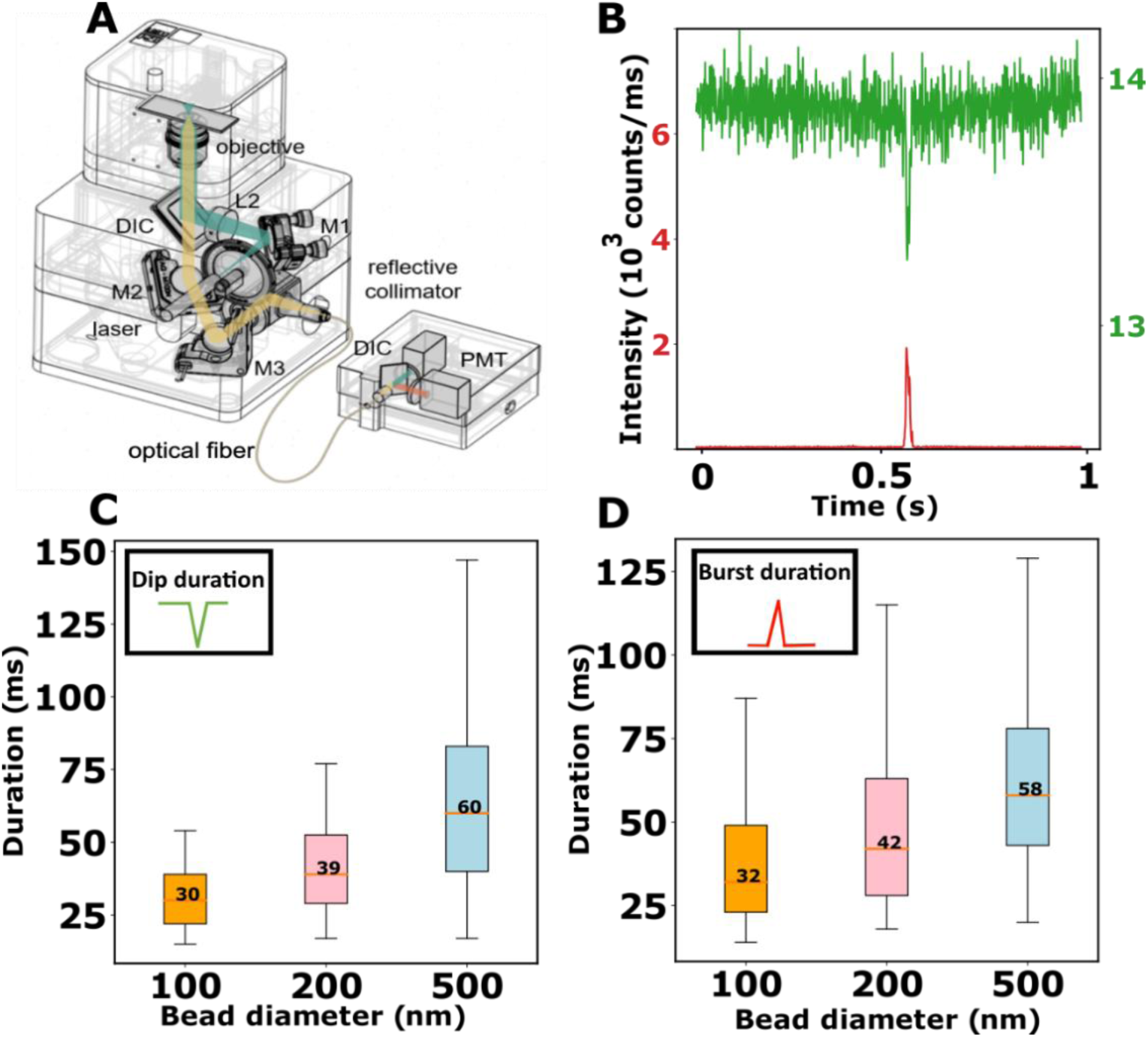
Detection of dye-labeled particles using a 3D-printed setup. **A**. Brick-MIC model A1 confocal microscope, developed by the Cordes lab^18^ and implemented in the flow virometer assay **B.** Coincident burst-dip pairs. Signal from a measurement of 200 nm diameter dye-labeled red beads in a 1 s time window (similar examples in Fig. S5). **C.** The durations of signal dips from coincident dip-burst detection events of dye-labelled polystyrene beads of diameter 100, 200 and 500 nm (accumulation of 37, 114 and 169 detection events from 4, 5 and 5 repetitions of 5 min. acquisitions, respectively, see Fig. S6) The durations are similar for coincident burst-dip pairs (P-values in Table S1). **D**. The duration of bursts in the specific signal formed from red-labeled beads of diameter 100, 200 and 500 nm (accumulation of 37 detection events from 4 repetitions of 5 min. acquisitions respectively, see Fig. S6). C,D measurement conduct in the presence of 500 µM fluorescein at a constant flow rate of 750 nL/min (2.08x10^-6^ m/s calculated mean velocity in microfluidic channel, see Eq. S1). P-values of comparisons between samples of different beads sizes in Table S1. For the histogram of durations of bursts solely from coincident burst-dip detection events, see Fig. S7. The orange horizontal line represents the median value.

Since a detection scheme based on one nonspecific signal does not reveal the identity of a particle, just its volume, we combined the information from the nonspecific signal with a *specific signal*, i.e., from spectrally distinct fluorophores that interact specifically with an epitope on the particle’s surface. The latter produces a fluorescence *burst*, as a temporal change in the specific signal (Fig. 1B, C). Both the changes in the nonspecific and specific signals are detected simultaneously using two spectrally-separated PMTs (Fig. 3A, B). Based on this layout, the detection of coincident pairs of nonspecific and specific signal changes, i.e., a coincident pair of dip and burst allows the unequivocal identification of a specific small-scale particle (Fig. 3B).

To evaluate this approach, we first used labeled polystyrene beads, where the red dyes on the beads mimic specific binders, such as antibodies (Figs. 1B, 3). Using this approach, it was possible to detect coincident bursts and dips (Fig. 3B) arising from beads with diameters ≥100 nm (Figs. 3C, S7). There is a well-established direct relationship between the duration of a signal burst and the diameter of a spherical small-scale particle ^22^ (Fig. 3D). The burst durations in the specific signal and the dip durations in the nonspecific signal are correlated (sample comparison P-values in Table S1). Therefore, one can use the duration of either the dip in the nonspecific signal or the burst in the specific signal to assess the particles’ diameters. Note, that only coincident dip-burst detection events are considered as specific particle events. It is noteworthy that analyzing the durations and sizes of both signals are setup-specific. Therefore, size assessment requires calibration with particle size standards (Fig. S8). The main differences that may occur between different experimental setups are due to differences in optical light path, type of objective lens, pinhole diameters, and laser powers and beam shapes.

Importantly, the size of a burst in the specific signal is affected by the number of dyes interacting with the observed particle. In contrast, the duration of a burst remains unaffected by the number of interacting dyes. Therefore, to ensure an unbiased assessment of small-scale particle diameters, we focus on analyzing burst durations of coincident signal events in both detection channels. In laminar flow, it is expected that spherical small-scale particles will experience increasing drag as their diameters increase. This, in turn, results in velocity reduction as their size increases, thereby extending burst durations.

Negative control measurements were performed with a mixture of unlabeled beads and free dyes to screen for the occurrence of coincidental events that do not occur due to a particle with both a relevant self-volume and a specific antibody interaction. Indeed, this control did not yield any detection (Table S2), hence the specificity of the detection scheme is maximal for the beads tested.

### Direct detection of Viruses

To further demonstrate the capabilities of our assay on biologically-relevant samples, we initially focused on detecting paraformaldehyde (PFA)-fixed and neutralized rVSV-ΔG-spike virus particle (Fig. 4A, B). The rVSV-ΔG-spike virus is a vesicular stomatitis virus (VSV), which is genetically-modified to exhibit the SARS-CoV-2 Spike protein, instead of its native G-protein^17^. In contrast to the observed dips in the nonspecific signal for polystyrene beads (Figs. 1B, 3), virus solutions showed bursts in the nonspecific signal, likely due to integration of free dyes onto the virus (Figs. 1C, 4A,C,E). Following this observation, we tested lower concentrations of free dye as well as fluorescein-labeled BSA (*Methods*). The aim was to minimize the potential interaction of free dyes with the interior of the virus, harnessing the inert characteristics of BSA^23^. We found that the bursts in the nonspecific signal could be distinguished, yet we observed spectral crosstalk between the nonspecific and specific signals, leading to coincidental false positive events. Careful adjustment of the BSA-fluorescein concentration enabled us to minimize these false positive events (Table S3, bold). All measurements using the Brick-MIC system with virus particles were conducted with 5 min. acquisition times, and for each measurement we calculated the detection event rate.

**Fig. 4.**
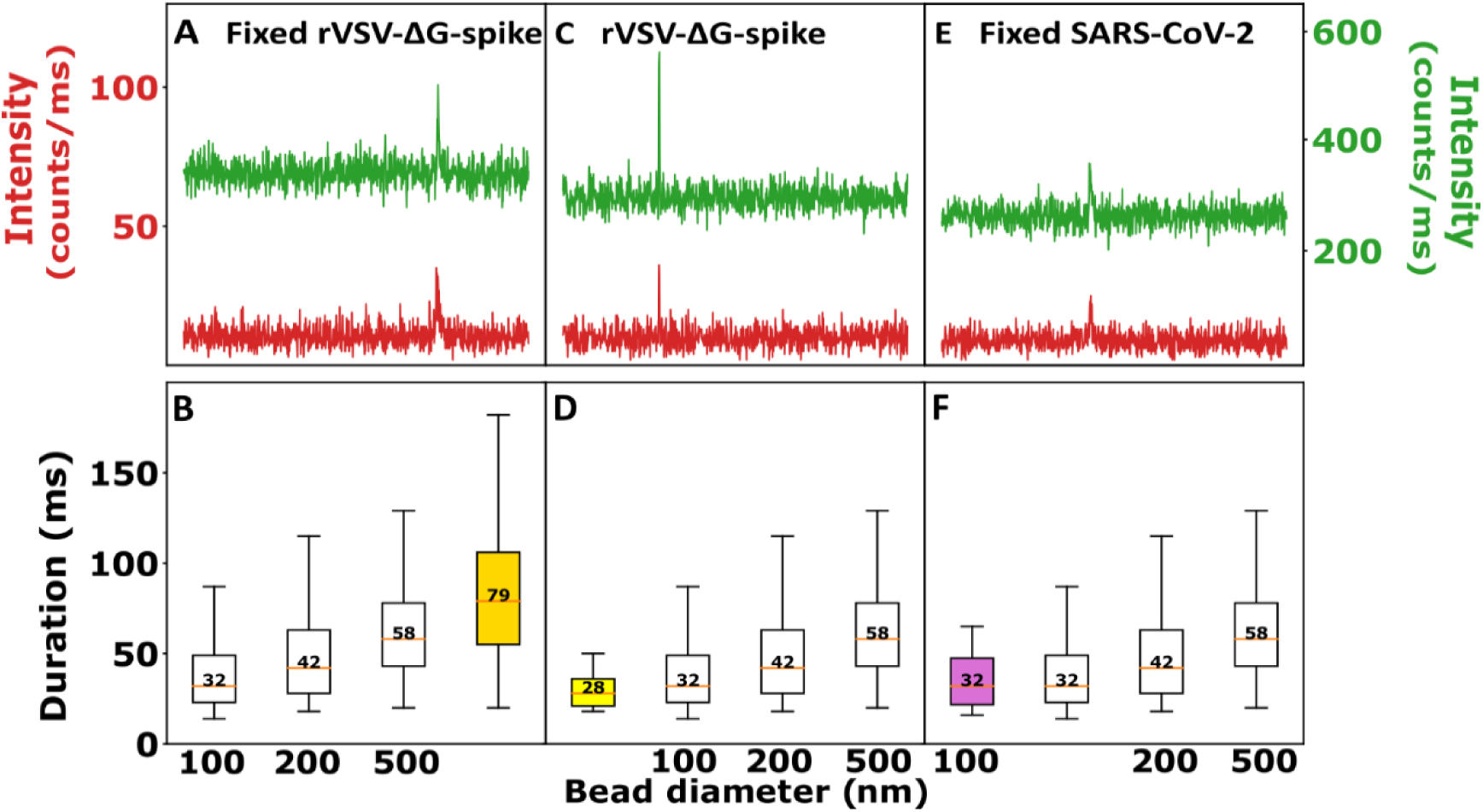
Counting single virus detection events. **A,B,C** 1 s data windows show coincident bursts, where the green bursts arise from the high-density accumulation of fluorescein on top of the particles detected in the nonspecific signal, and the red bursts arise from fluorescence of the bound antibodies observed in the specific signal. Similar examples in Fig. S9. **B,D,F** The duration of bursts in the specific signal of coincident burst events of detected viruses shown together with the burst durations of beads with different diameters. All virus samples were measured in the presence of eq. 10 µM BSA-fluorescein at a constant flow rate of 750 nL/min (2.08x10^-6^ m/s calculated mean velocity in microfluidic channel, see Eq. S1) (**B**) fixed rVSV-ΔG-spike (**D**) live rVSV-ΔG-spike (**F**) fixed SARS-CoV-2 (accumulation of detection events from repetitions and biological repeats, see Fig. S10). The orange horizontal line represents the median value. P-values of comparisons between samples of different beads sizes in Table S1.

For specific detection of rVSV-ΔG-spike, we used red dye-labeled fluorescent antibodies targeting the SARS-CoV-2 spike protein as the source of bursts in the specific signal and BSA-fluorescein as the source of changes in the nonspecific signal. This approach facilitated the identification of antibody-labeled virus particles via coincident bursts in both the nonspecific and specific signals (Fig. 4). The rVSV-ΔG-spike virus is bullet-shaped with a diameter of ∼70 nm and height of ∼180 nm^24^. Since the dependence of burst durations on particle volumes was established using spherical beads (Fig. 3D), the volume of a rVSV-ΔG-spike virus corresponds to that of a spherical particle with a diameter ∼100 nm, and indeed the experimental results indicate this similarity (Fig. 4C, D; P-values in Table S1). In contrast, the durations of signal bursts of PFA-fixed rVSV-ΔG-spike viruses, which tend to cluster^25^, correspond to spherical particleswith diameters of >500 nm (Fig. 4A, B; P-values in Table S1). These results clearly indicate a deviation from detection of non-clustered single particles.

To verify the detection specificity of our approach for rVSV-ΔG-spike viruses, we performed a control measurement of nonspecific antibodies with rVSV-ΔG-spike viruses (0.10±0.10, standard error of the mean, SEM, detections per 5 min. acquisitions, Table S4), which should not bind. In addition, background measurements of an irrelevant protein, the human serum albumin (HSA), were conducted with and without specific dye-labeled antibodies (0.17±0.09 and 0.14±0.09, SEM, detections per 5 min. acquisitions, respectively, Table S4). In measurements of viruses in the presence of specific antibodies (Table S4, condition F, bolded), an average of 8.72±1.02 (SEM) specific detection events are reported within 5 min. acquisition times. We estimate the false detection rate (FDR) to be 2.0±1.4% (SEM), based on the results of measuring fixed rVSV-ΔG-spike viruses (FDR calculation in Eqs. S2, S3).

The ability to count single virus detection events with specific properties was further confirmed with fixed and neutralized SARS-CoV-2 viruses (Fig. 4E, F). These viruses are known to be spherical with diameters in the 60-140 nm range^26^. In agreement, we observe burst parameters that significantly differ from those of beads with diameters ≥200 nm. However, these observations do not significantly differ from beads with 100 nm diameters (Fig. 4E, F; P-values in Table S1). The concentration of the PFA-fixed SARS-CoV-2 virus particles was lower than the PFA-fixed rVSV-ΔG-spike particles (1x10^5^/mL versus 1x10^8^/mL, respectively). Since the level of clustering depends on overall concentration, the PFA-fixed SARS-CoV-2 particles cluster less than the PFA-fixed-rVSV-ΔG-spike particles in our measurements.

### Increasing sensitivity for detection of nanoparticles at lower concentrations

While the presented results validate the detection specificity, further characterization of the sensitivity of the assay is required to determine the minimal required particle concentrations or viral loads within a given acquisition time. In the results of further experiments (Fig. 4) the SARS-CoV-2 loads were lower than those of the rVSV-ΔG-spike, i.e., 10^5^ versus 10^8^ particles/mL, respectively. These loads further decreased during the PFA neutralization process, which facilitate using SARS-CoV-2 viruses in labs with biosafety level 1^25^. Out of these reasons, the acquisition time was significantly longer for the SARS-CoV-2 viruses (Fig. S10).

We aspire to detect at least 30 detection events, within the shortest possible acquisition time, and we consider ∼1 detection event at most to be a false event, based on the FDR estimate in the range 0.6-3.4% (Table S4). We refer to the lowest viral particle load and measurement conditions that will lead to such results as the *sensitivity limit*. Within the current results (Fig, 4), the required acquisition time would be ∼20 min., however for particle loads in the range 10^7^-10^8^ particles/mL. To facilitate detection of viruses from patient samples out of bodily fluids it is required to adapt the assay for loads in the range 10^3^-10^7^ particles/mL, which is the typical SARS-CoV-2 range of loads in saliva^27^. Doubling the flow rate to 1,500 nL/min (4.16x10^-6^ m/s), moderately increases the mean detection rate to 23.6±3.8 (SEM) particles per 5 min. acquisition time (Fig. S11). This, however, results in shorter burst durations reducing the detectability of signal changes (Fig. S11). Overall, flow rate changes using the same fluidics do not enhance the sensitivity sufficiently, i.e., by orders of magnitudes.

To assess the limits of our sensitivity, we implemented microfluidic hydrodynamic focusing using polystyrene beads and our laboratory confocal setup (Fig. 5A) just as a proof of concept. Using a commercial 3-to-1 microfluidic chip (Fig. 5A), we achieved microfluidic hydrodynamic focusing, as can be seen from the width of the analyte stream (Fig. 5B). Indeed, by increasing the sheath flow rate relative to a constant analyte flow rate, the cross-section area of the analyte stream decreases (Fig. 5F) relative to the constant cross-section area of the probe volume (i.e. the confocal volume) governed by the objective lens^28^. After focusing the probe volume within the hydrodynamically-focused analyte stream (Fig. 5A, red dot) we counted single detection events with fluorescent beads (Fig. 5C). Using this approach, the count rate of detection events was increased from ∼5 to ∼150 detections per 5 min. acquisition time for an increase of the sheath flow rate from 2 to 30 μL/min (1.39x10^-6^ to 2.08x10^-5^ m/s), respectively, using a constant analyte flow rate of 1 μL/min (6.93x10^-7^ m/s). The maximal focusing of the analyte stream, from an analyte stream width of 1,000 μm down to 30 μm, was achieved at a sheath flow rate of 700 μL/min (4.85x10^-4^ m/s; Fig. 5F).

**Fig. 5.**
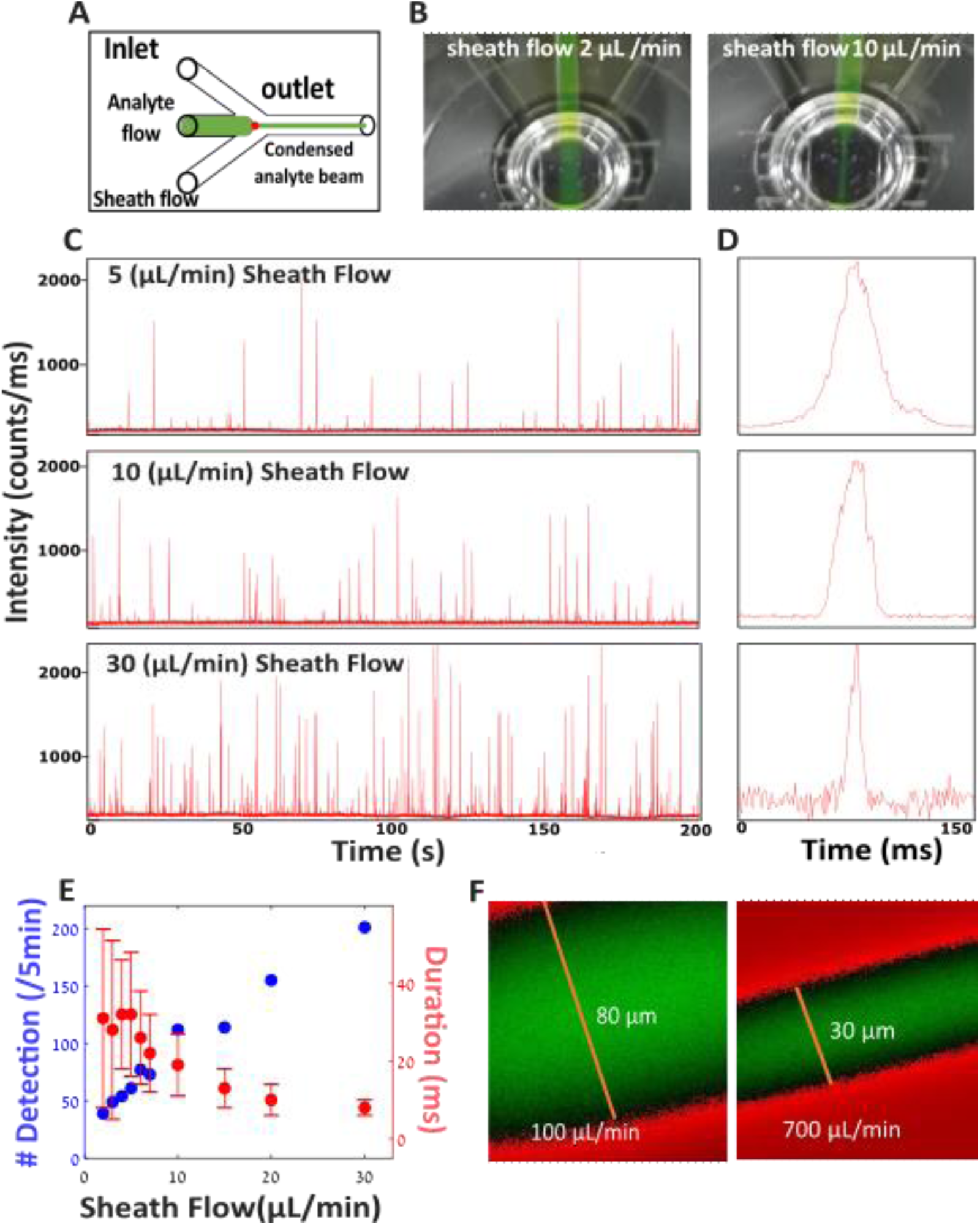
Increasing particle detection rate with microfluidic hydrodynamic focusing. **A** Concept illustration. Analyte stream (green) is focused at the junction between analyte flow channel (central channel) and two peripheral sheath flow channels (lower and upper channels) at higher flow rates relative to the analyte flow rate. Probe volume (red dot) placed at center of hydrodynamically-focused analyte. **B** Analyte flow rate of 1 μL/min (6.93x10^-7^ m/s calculated mean velocity in analyte inlet, see Eq. S1) is kept straight or focused as the sheath flow increases from 2 μL/min (1.39x10^-6^ m/s; left) to 10 μL/min (6.93x10^-6^ m/s; right). One can observe narrowing of the fluorescein analyte flow. **C** Signal acquisition from dye-labeled 1,000 nm diameter latex beads flowing as analyte at a rate of 1 µL/min before the hydrodynamic focusing junction (red bursts). The sheath flow rate was gradually increased from 5 µL/min (3.46x10^-6^ m/s; top) to 30 µL/min (2.08x10^-5^ m/s; bottom), while the analyte flow rate was kept steady. **D** Representative burst. One can observe shortening of burst durations as sheath to analyte flow rate ratio increases. **E** The average amount of detected single particle bursts (blue) and their durations (red) as a function of sheath to analyte flow rate ratio. **F** The higher the sheath to analyte flow rate ratio is, the more focused does the analyte stream become. Sheath flow rates 100 (left) and 700 (right) µL/min (6.93x10^-5^ and 4.85x10^-4^ m/s, respectively) and 1 µL/min analyte flow rate lead to decreasing width of the analyte stream to 80 and 30 μm, respectively.

Using beads in laminar flow in a one-inlet-one-outlet microfluidic channel, the detection sensitivity limit was achieved for a load of 10^8^ particles/mL (Figs. 2, 3). By extrapolation, the hydrodynamic focusing approach (Fig. 5) can assist in reaching this sensitivity limit with loads as low as 3x10^6^ particles/mL (i.e., 30 µm / 1,000 µm x 10^8^ particles/mL). We note that we already used the one-inlet-one-outlet microfluidic channel with a 100x1,000 µm^2^ cross-section area to count virus detection events for virus samples at loads of 10^5^ particles/mL. Therefore, by the same extrapolation, we speculate that the hydrodynamic focusing approach can, in principle, assist in reaching the sensitivity limit for these viruses with loads as low as 3x10^3^ particles/mL for acquisition times >15 min. This value assists in covering the biomedically-relevant range of viral loads^27^.

## Discussion

We developed a virus detection assay that functions as a sensitive and specific small-scale particle counter, relying on confocal-based detection in combination with microfluidic laminar flow. We demonstrated the use of this assay in the rapid and accurate detection of >100 nm particle diameters, such as polystyrene beads and on virus particles containing the SARS-CoV-2 S protein on their surface. Importantly, the method allows the detection of low particle loads based on non-amplifiable targets, such as proteins, via the interaction with dye-labeled antibodies, in contrast to amplified signal-based methods, such as PCR or ELISA. This advantage could reduce potential false negative events in cases of too many amplification cycles, which could propagate errors^29,30^. While ELISA is the method of choice for identifying protein targets using antibodies, it has longer acquisition times, as well as higher sample consumption relative to the implementation of our assay^31,32^.

In addition, we demonstrated how the sensitivity of the assay can be further improved by microfluidic hydrodynamic focusing. Yet, the improvement demonstrated was achieved using our laboratory confocal setup with polystyrene beads. The improved detection sensitivity has not yet been achieved on the portable 3D-printed setup, due to the current lack of control over the *x* and *y* positioning of the sample. The follow-up prototype of the 3D-printed setup will include such control over sample position. It will allow proper positioning of the probe volume relative to the hydrodynamically-focused analyte stream (Fig. 5F). Additionally, improved microfluidics layout (e.g., T-shaped junction instead of the current Y-shaped junction^28^) and pumps with higher sensitivities (e.g., perfusion pumps) can be combined to increase the detection sensitivity and theoretically reach the sensitivity limit with even lower loads.

Besides technical improvements, we envision an extension of the assay towards using additional spectrally-resolved detections of additional antibodies and spectral multiplexing. The potential use of three detection channels instead of two greatly extends the possibilities to use additional fluorescent signals, such as the coincident detection of more than one antigen or binding target per particle, or of different types of particles, such as antibodies against different antigens of the same virus or for two different viruses (Fig. S12).

In this work, we demonstrated the detection based on antibody-antigen interactions. However, the identification can also be based on additional specific interactions against specific markers of the target such as nucleic acids via intercalating dyes or hybridization interactions^33^. This technology can also be adjusted for detecting other bio-nanoparticles carrying specific antigens that might be of biomedical importance or mixtures, especially if these particles do not contain specific biomolecules that can be amplified. This solution can provide a platform for the detection of specific targets at scarce amounts in bodily fluids, e.g., exosomes and biomarkers for different cancer types^34^ or perhaps even for early detection of neurodegenerative diseases-related exosomes^35^.

In summary, the specific and sensitive counting of bio-NPs one-at-a-time, within minutes, using a simple assay and an affordable experimental setup could be considered for assisting in efficient diagnostics alongside existing traditional techniques.

## Methods & Materials

Unless stated otherwise, all chemical compounds were purchased from Sigma and used as received.

### Polystyrene beads measurement preparation

Polystyrene beads with diameters of 1,100 nm, 600 nm, and 300 nm, along with red polystyrene (FluoSpheres Size Kit #1, carboxylate-modified, red, with an excitation/emission λmax of 580/610 nm, 2% solid, from Thermo-Fisher, having diameters of 100 nm, 200 nm, and 500 nm) were added to a buffer. The buffer contain75 mM KCl, 0.025% Triton, 20 mM TRIS pH 8.6 and 1% PEG 10,000 to a particle concentration ranging from 10^15^ to10^8^ particles/mL. The mixture underwent sonication for 20 to 35 minutes and then free dye (Fluorescein, excitation/emission λ_max_= 488/510 nm) was added to the final concentration of 500 µM. The prepared sample was then loaded into a 1 mL syringe for subsequent use in microfluidics.

### Viruses

rVSV-ΔG-spike, a replication competent recombinant rVSV-ΔG-spike virus, in which the glycoprotein (G) of VSV was replaced by the Spike protein of SARS-CoV-2 (IIBR, Israel). Before use, virus stocks (1x10^8^/mL) were titred on Vero E6 cells using Plaque forming unit assay (PFU) as previously described^17^. In brief, Vero E6 cells were seeded (5x10^5^ cells/12 well plates) and grown overnight. Dilutions of viruses were prepared and used for infecting Vero E6 monolayers. Plates were incubated for 1 hour at 37 °C in order to allow viral adsorption. Then, 1 mL/well of overlay media (0.4% Tragacanth, Merck) was added to each well and plates were incubated at 37°C, 5% CO_2_ for 48 hours. The media was then aspirated and the cells were fixed and stained with 500 µL/well of crystal violet solution (Biological Industries). The number of plaques in each well was determined and rVSV-ΔG-spike titer was calculated.

SARS-CoV-2 viruses (GISAID accession EPI_ISL_406862; the IMB, Germany) Virus stocks (1x10^5^/mL) were propagated (four passages) and tittered on Vero E6 cells before use, as previously described^17^,. Handling and working with SARS-CoV-2 virus were conducted in a BSL3 facility in accordance with the biosafety guidelines of the IIBR. SARS-CoV-2 sample was neutralized and fixed using 14% paraformaldehyde (PFA).

### Virus fixation and neutralization

To allow work under BSL1 conditions, virus preparations of SARS-CoV-2 and rVSV-ΔG-spike were neutralized and fixed with 14% PFA in PBS for 20 min. PFA was removed by dialysis using cellulose acetate membrane (100 kDa molecular weight cutoff (Harvard Apparatus) at room temperature for 4 hours incubation against 4 L of PBS. At least four additional dialysis cycles were applied with fresh PBS before a final cycle of overnight incubation at 4°C.

### Antibody fluorescent labeling

Anti-spike antibodies (5 µg/µL rabbit IgG polyclonal anti-receptor binding domain, RBD, of the viral spike protein; IIBR, Israel)^17^ were used for the specific detection of both SARS-CoV-2 and rVSV-ΔG-spike viruses. Antibodies (100 µg) were labeled with Alexa Fluor® 594 Protein Labeling Kit (Thermo-Fisher Scientific) according to the manufacturer’s instructions.

### Virus measurement preparation

The SARS-CoV-2 and rVSV-ΔG-spike viruses were mixed with dye-labeled rabbit anti-spike antibodies (excitation λ_max_ = 594 nm, IIBR, Israel, see Antibody fluorescent labeling)^17^ to reach a final concentration of 1 µg/mL. The mixture was incubated while rotating for 30 minutes in the dark at room temperature. As a control, we measured viruses without antibodies or with nonspecific antibodies (Goat Anti-Rabbit IgG H&L Alexa Fluor® 594). BSA-fluorescein was added to achieve a final concentration equivalent to 10 µM fluorescein, based on fluorescein absorption and fluorescence readout (∼300 counts per s). Subsequently, the sample was loaded into a 1 mL syringe.

### Laboratory confocal-based setup

The confocal-based setup (ISS™, USA) is assembled on top of an Olympus IX73 inverted microscope stand. We use a pulsed picosecond fiber laser for excitation (λ = 532 nm, pulse width of 100 ps FWHM, operating at 20 MHz repetition rate with 150 μW laser power measured at the back aperture of the objective lens). The laser beam passes through a polarization-maintaining optical fiber and is then further shaped by a quarter-wave plate and a linear polarizer. A dichroic beam splitter with high reflectivity at 532 nm (ZT532/640rpc, Chroma, USA) reflects the light through the optical path to a high numerical aperture (NA) super Apo-chromatic objective (UPLSAPO100XO, 100X, 1.4 NA, oil immersion, Olympus, Japan), which focuses the light onto a small confocal volume, referred to in this work as the effective excitation volume. The microscope collects the fluorescence from the excited molecules through the same objective and focuses it with an achromatic lens (f = 100 mm) onto a 100 μm diameter pinhole (variable pinhole, motorized, tunable from 20 μm to 1 mm), and then re-collimates it with an achromatic lens (f =100 mm). Fluorescence was filtered from other light sources (transmitted scattering) with a 510/20 nm band-pass filter (FF01-510/20-25, Semrock Rochester NY, USA) and detected using a hybrid photomultiplier (Model R10467U-40, Hamamatsu, Japan), routed through a CFD unit (ISS^TM^, USA) and a correlator (ISS^TM^, USA) to the acquisition software. It is noteworthy that the nonspecific signal arising from the fluorescence of free fluorescein dyes, was measured following excitation at the red edge of fluorescein absorption spectrum with a 532 nm laser and its fluorescence was collected in its maximal fluorescence range using a 510/20 nm filter. We perform data acquisition using the VistaVision software (version 4.2.095, 64-bit, ISS™, USA). We combine the described system with a microfluidic channel (µ-Slide VI 0.5 Glass Bottom, Ibidi and µ-Slide III 3in1, Ibidi for hydrodynamic focusing) and a fine syringe pump (single channel multi-mode 2 syringe pump, MRClab) to achieve microfluidic laminar flow, while focusing the probe volume of the focused laser at the center of the channel (Fig. 1D).

### 3D-Printed portable setup

The 3D-printed setup shown in figure 3A was designed as described previously^18^. The setup uses a 532 nm wavelength CW laser diode (5 mW output; CPS532, Thorlabs) as excitation light source. The beam is attenuated by a neutral density filter (OD =1.5) and expanded with a telescope consisting of a bi-concave (f = -50 mm, KBC043AR.14, Newport) and a plano-convex lens (f = 150 mm, LA1433-A-ML, Thorlabs). A dichroic beam splitter with high reflectivity at 532 nm (ZT532/640rpc, Chroma, USA) separates excitation and emission beams into and from a high numerical aperture (NA) apo-chromatic objective (100X, NA=1.4, oil immersion, Olympus, Japan). The emitted fluorescence is collected by the same objective and directed via a mirror in a piezo directed optical mount (AG-M100N, Newport) through an inversely mounted 12 mm reflective collimator (RC12FC-F01, Thorlabs), which focuses and couples the emission beam into a multimode optical fiber (10 µm fiber core diameter, M64L01, Thorlabs). The fiber directs the coupled emission light into a detection box, where it is collimated with a fixed focus collimator (F220FC-532, Thorlabs) and then spectrally split into two separate photon streams by a dichroic mirror (ZT640rdc longpass, Chroma, USA). Individual photon streams are filtered with bandpass filters (for the unspecific channel: FF01-510/20-25, Semrock Rochester NY, USA; for the specific channel: ET700/75m, Chroma) and detected by two distinct photomultiplier tubes (for the unspecific channel: H10682-210, Hamamatsu, Japan; for the specific channel: H10682-01, Hamamatsu, Japan). The detector outputs were recorded by a counter/timer device module (USB-CTR04, Measurement Computing, USA). Much like in the laboratory confocal-based setup, also here, the nonspecific signal arising from the fluorescence of free fluorescein dyes, was measured in anti-Stokes mode. We performed data acquisition using a custom-made acquisition software written in Python that is freely available via https://github.com/klockeph/mcc-daq-acquisition. Similar to the use in the confocal-based setup (see above), we combine the described system with a microfluidic channel (µ-Slide VI 0.5 Glass Bottom, Ibidi) and a fine syringe pump (single channel multi-mode 2 syringe pump, MRClab) to achieve microfluidic laminar flow, while focusing the probe volume of the focused laser at the center of the channel (Fig. 1D).

### Microfluidic slides used

To achieve laminar flow, we conduct the experiment using a commercially available microfluidic channel and a syringe pump. For validation of our assay and for the detection of the virus we use a one-inlet-one-outlet microfluidic channel (Ibidi µ-slide VI 0.1 cannel, with cross-section of 100x1,000 µm^2^). For the hydromantic focusing we use a three-inlet-one-outlet microfluidic slide (µ-Slide III 3in1, 3-channels with cross-section 400x1,000 µm^2^ combined to a 1-channel with cross-section 400x3,000 µm^2^). To convert the flow rate to attain the calculated mean velocity in the microfluidic channel, use the relation shown in the SI (Eq. S1).

### Data analysis

All data analyses were performed using Python-written code (see https://doi.org/10.5281/zenodo.10277721). Data acquisition from both setups were performed after data conversion to 1 ms time bin traces. The principle of finding a dip/burst in the specific or nonspecific signal (depending on what we measured; see Fig. 1) is based on finding the local extremum of the signal and define the initial and final time points of the event. First, the data is divided to 1,000 ms time frames. Then, the data is smoothed using the Savitzky-Golay algorithm (*scipy.signal.savgol_filter*; x,25,3). For each time frame the mean and the standard deviation is calculated. We define a threshold line for each data frame as the value of 3-5 standard deviations above/below the mean (see Table S5) to find potential burst/dip (Fig. S13, green line). Then the local extremum point (Fig. S13, yellow dot), that is between two intersecting time points (Fig. S13, green dots), between the data (Fig. S13, blue line) and threshold line (Fig. S13, green line; found using *numpy.argwhere*), are identified. In the next step, the initial and final time points (Fig. S13, red dots) of the dip/burst are found by searching for the intersection time point between the data and line equal to the mean (Fig. S13, orange line), before and after the local extremum point (Fig. S13). The signal size is calculated by subtracting the mean value from the value of the extremum point. The signal duration is calculated by subtracting the value of the final time point from the value of the initial time point. For coincident detection events, in each data frame, the initial and final time points of the event (dip or burst) in the nonspecific signal was compared with the initial and final time points of the burst in the specific signal to find if there is a time overlap between the two. The data from all the correlated signals from all the data frames of all the measurement are accumulate.

## Acknowledgments

The authors would like to thank the Israel Institute for biological Research (IIBR): (1) Dr. Tomer Israeli and Dr. Hadas Tamir of the department of infectious diseases for providing the SARS-CoV-2, (2) the department of biotechnology for providing the rVSV-ΔG-spike, and (3) Dr. Efi Makdasi of the department of biochemistry and molecular genetics for the anti-Spike antibodies. This project was supported by the Israel science foundation (ISF; grants 556/22 to E.L. and 3565/20 to E.Z. and E.L., within the KillCorona – Curbing Coronavirus Research Program), the national institutes of health (grant R01 GM130942 to E.L. as subaward), the Bundesministerium für Bildung und Forschung (KMU grant „quantumFRET“ to T.C.), the European Comission (ERC-STG 638536 – SM-IMPORT to T.C.) and by the Milner Fund (to E.L.).

## Author contributions

E.L. and T.C. initiated the project. P.D. and E.L. designed and conceived the study and the experiments. P.D., Y.R, O.M. and R.A. prepared samples, G.G.M.M. built the 3D-printed microscopy platform, P.K design the acquisition software for the 3D-printed microscopy platform P.D. and Y.R. conducted experiments, P.D. analyzed data, P.D. and Y.R prepared figures. E.Z., T.C. and E.L. acquired funding. E.L. supervised the study. P.D. and E.L. wrote the initial draft of the manuscript, which was reviewed, edited and approved by all authors.

## Additional information

G.G.M.M., P.D., Y.R., T.C. and E.L. have submitted a patent about the flow-based detection scheme presented in this work: Flow virometer for rapid detection of intact viruses. (2022), PCT Publication No. WO2022172208A1. G.G.M.M., P.D., Y.R., T.C. and E.L. declare commercial interest in this patent.

## Notes

https://doi.org/10.5281/zenodo.10277721

### Summary of Updates

The text and one figure have been revised.

## References

1. Ciotti, M. et al. The COVID-19 pandemic. Crit. Rev. Clin. Lab. Sci. 57, 365–388 (2020).

2. Chartrand, C., Tremblay, N., Renaud, C. & Papenburg, J. Diagnostic Accuracy of Rapid Antigen Detection Tests for Respiratory Syncytial Virus Infection: Systematic Review and Meta-analysis. J. Clin. Microbiol. 53, 3738–3749 (2015).

3. Grandien, M. Viral diagnosis by antigen detection techniques. Clin. Diagn. Virol. 5, 81–90 (1996).

4. Eshghifar, N., Busheri, A., Shrestha, R. & Beqaj, S. Evaluation of Analytical Performance of Seven Rapid Antigen Detection Kits for Detection of SARS-CoV-2 Virus. Int. J. Gen. Med. Volume 14, 435–440 (2021).

5. Klein, D. Quantification using real-time PCR technology: applications and limitations. Trends Mol. Med. 8, 257–260 (2002).

6. Watzinger, F., Ebner, K. & Lion, T. Detection and monitoring of virus infections by real-time PCR. Mol. Aspects Med. 27, 254–298 (2006).

7. Mackay, I. M. Real-time PCR in virology. Nucleic Acids Res. 30, 1292–1305 (2002).

8. Andrew, A., et al. Diagnostic accuracy of serological tests for the diagnosis of Chikungunya virus infection: A systematic review and meta-analysis. PLoS Negl. Trop. Dis. 16, e0010152 (2022).

9. Tantuoyir, M. M. & Rezaei, N. Serological tests for COVID-19: Potential opportunities. Cell Biol. Int. 45, 740–748 (2021).

10. Lippé, R. Flow Virometry: a Powerful Tool To Functionally Characterize Viruses. J. Virol. 92, e01765–17 (2018).

11. Brussaard, C. P. D., Marie, D. & Bratbak, G. Flow cytometric detection of viruses. J. Virol. Methods 85, 175–182 (2000).

12. Stoffel, C. L., Kathy, R. F. & Rowlen, K. L. Design and characterization of a compact dual channel virus counter. Cytometry A 65A, 140–147 (2005).

13. Zamora, J. L. R. & Aguilar, H. C. Flow virometry as a tool to study viruses. Methods 134–135, 87–97 (2018).

14. Niu, Q., et al. Quantitative Assessment of the Physical Virus Titer and Purity by Ultrasensitive Flow Virometry. Angew. Chem. Int. Ed. 60, 9351–9356 (2021).

15. Hepp, C. et al. Viral detection and identification in 20 min by rapid single-particle fluorescence in-situ hybridization of viral RNA. Sci. Rep. 11, 19579 (2021).

16. Shiaelis, N. et al. Virus Detection and Identification in Minutes Using Single-Particle Imaging and Deep Learning. ACS Nano 17, 697–710 (2023).

17. Yahalom-Ronen, Y. et al. A single dose of recombinant VSV-ΔG-spike vaccine provides protection against SARS-CoV-2 challenge. Nat. Commun. 11, 6402 (2020).

18. Moya Munoz, G. G., et al. Single-molecule detection and super-resolution imaging with a portable and adaptable 3D-printed microscopy platform *(Brick-MIC)*. http://biorxiv.org/lookup/doi/10.1101/2023.12.29.573596 (2023) doi:10.1101/2023.12.29.573596.

19. Drori, P., Razvag, Y., Moya, G., Cordes, T. & Lerner, E. Flow virometer for rapid detection of intact viruses.

20. Wennmalm, S. & Widengren, J. Inverse-Fluorescence Cross-Correlation Spectroscopy. Anal. Chem. 82, 5646–5651 (2010).

21. Wennmalm, S. Inverse-fluorescence correlation spectroscopy more information and less labeling. Front. Biosci. S3, 385–392 (2011).

22. Edel, J. B. & De Mello, A. J. Single Particle Confocal Fluorescence Spectroscopy in Microchannels: Dependence of Burst Width and Burst Area Distributions on Particle Size and Flow Rate. Anal. Sci. 19, 1065–1069 (2003).

23. Sweryda-Krawiec, B., Devaraj, H., Jacob, G. & Hickman, J. J. A New Interpretation of Serum Albumin Surface Passivation. Langmuir 20, 2054–2056 (2004).

24. Rozo-Lopez, P., Drolet, B. & Londoño-Renteria, B. Vesicular Stomatitis Virus Transmission: A Comparison of Incriminated Vectors. Insects 9, 190 (2018).

25. Möller, L. et al. Evaluation of Virus Inactivation by Formaldehyde to Enhance Biosafety of Diagnostic Electron Microscopy. Viruses 7, 666–679 (2015).

26. Zhu, N. et al. A Novel Coronavirus from Patients with Pneumonia in China, 2019. N. Engl. J. Med. 382, 727–733 (2020).

27. To, K. K.-W. et al. Temporal profiles of viral load in posterior oropharyngeal saliva samples and serum antibody responses during infection by SARS-CoV-2: an observational cohort study. Lancet Infect. Dis. 20, 565–574 (2020).

28. Hong, S. et al. Microfluidic three-dimensional hydrodynamic flow focusing for the rapid protein concentration analysis. Biomicrofluidics 6, 024132 (2012).

29. Yang, S. & Rothman, R. E. PCR-based diagnostics for infectious diseases: uses, limitations, and future applications in acute-care settings. Lancet Infect. Dis. 4, 337–348 (2004).

30. Buchta, C. et al. Variability of cycle threshold values in an external quality assessment scheme for detection of the SARS-CoV-2 virus genome by RT-PCR. Clin. Chem. Lab. Med. CCLM 59, 987–994 (2021).

31. Caygill, R. L., Blair, G. E. & Millner, P. A. A review on viral biosensors to detect human pathogens. Anal. Chim. Acta 681, 8–15 (2010).

32. Aydin, S. A short history, principles, and types of ELISA, and our laboratory experience with peptide/protein analyses using ELISA. Peptides 72, 4–15 (2015).

33. Ha, T., Kaiser, C., Myong, S., Wu, B. & Xiao, J. Next generation single-molecule techniques: Imaging, labeling, and manipulation in vitro and in cellulo. Mol. Cell 82, 304–314 (2022).

34. Jiang, L., Gu, Y., Du, Y. & Liu, J. Exosomes: Diagnostic Biomarkers and Therapeutic Delivery Vehicles for Cancer. Mol. Pharm. 16, 3333–3349 (2019).

35. Hornung, S., Dutta, S. & Bitan, G. CNS-Derived Blood Exosomes as a Promising Source of Biomarkers: Opportunities and Challenges. Front. Mol. Neurosci. 13, 38 (2020).

