## Supplementary Information for "Rapid and specific detection of single nanoparticles and viruses in microfluidic laminar flow via confocal fluorescence microscopy"

[3] Physical and Synthetic Biology. Faculty of Biology, Ludwig-Maximilians-Universität München, Großhadernerstr. 2-4, 82152, Planegg-Martinsried, Germany.

[4] The Center for Nanoscience and Nanotechnology, The Hebrew University of Jerusalem, Jerusalem 9190401, Israel.

**Supplementary Text**

To convert the flow rate from nL/min to m/s we use the following relation (Eq. S1)

$$v\ \left[\frac{m}{s}\right] = \frac{V/t\left[\frac{nL}{min}\right]}{w[\mu m]\cdot h[\mu m]}\cdot\frac{1x10^{-6}\left[\frac{mL}{nL}\right]}{1x10^{-8}\left[\frac{cm^2}{\mu m^2}\right]}\cdot\frac{1}{60}\left[\frac{s}{min}\right]\cdot\frac{1}{100}\left[\frac{m}{cm}\right] = 2.77x10^{-4}\cdot\frac{V/t\left[\frac{nL}{min}\right]}{w[\mu m]\cdot h[\mu m]} \quad \text{(Eq. S1)}$$

where $t$ is time, $v$ is the velocity, $V$ is volume, $w$ is the width of the channel and $h$ is the height of the channel.

The false detection rate (FDR), *F* was calculated according to the following relation (Eq. S2)

$$F = \frac{A}{B} = \frac{0.17}{8.72} = 0.02 = 2\% \quad \text{(Eq. S2)}$$

where, *F* is the ratio between the mean number of events from a negative control, *A*, and the mean number of events in an actual measurement with both true positives and potential false positives, *B*. The uncertainty of the FDR, *dF*, is calculated according to the following relation (Eq. S3)

$$dF = \sqrt{\left(\frac{\partial F}{\partial A}dA\right)^2 + \left(\frac{\partial F}{\partial B}dB\right)^2} = \sqrt{\left(\frac{dA}{B}\right)^2 + \left(\frac{AdB}{B^2}\right)^2} = \sqrt{\left(\frac{0.095}{8.722}\right)^2 + \left(\frac{0.176x1.018}{8.722^2}\right)^2} = 0.014 = 1.4\% \quad \text{(Eq. S3)}$$

where *dA* and *dB* are the standard error of the mean (SEM) of *A* and *B*, from several repeats. The values added to Eqs. S2 and S3 are taken from Table S4, conditions B and F (conditions A-E are different types of negative control, and out of them B gave the maximal number of coincident events).

### Supplementary Figures

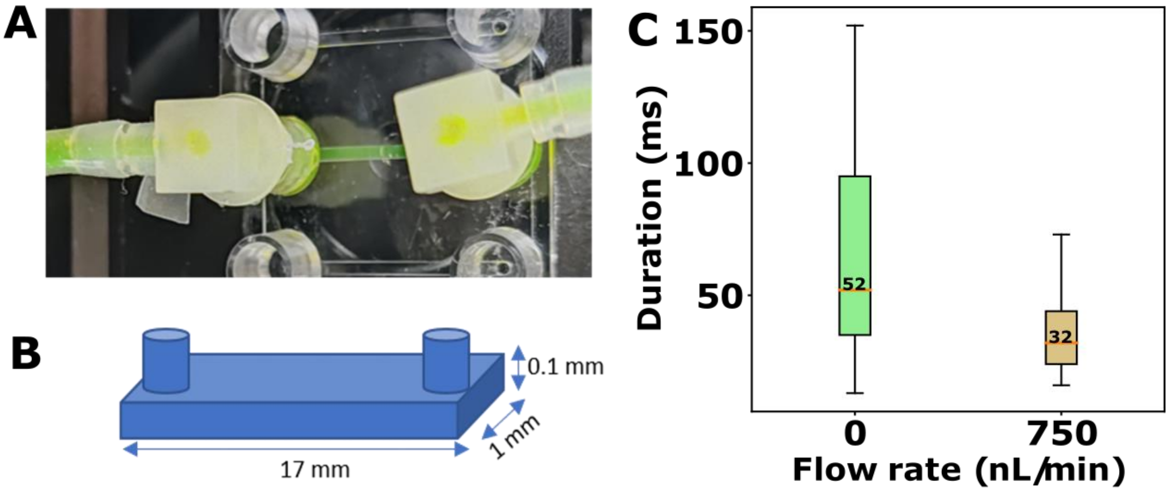


**Fig. S1. Microfluidic laminar flow measurements. A B**. Ibidi µ-slide VI 0.1, with sample flow through, mounted on top of microscope stage. **C**. Distributions of signal duration of 7 min. measurements of 300nm beads in the presence of 500 µM fluorescein with either without flow (green, accumulation of 65 detection events from 3 repetitions) or flow rate of 750 nL/min ($2.08 \times 10^{-6}$ m/s calculated mean velocity in microfluidic channel Eq. S1; brown, accumulation of 314 detection events from 4 repetitions) P-value=*** between sample (see Table S1). The orange horizontal line represents the median value.

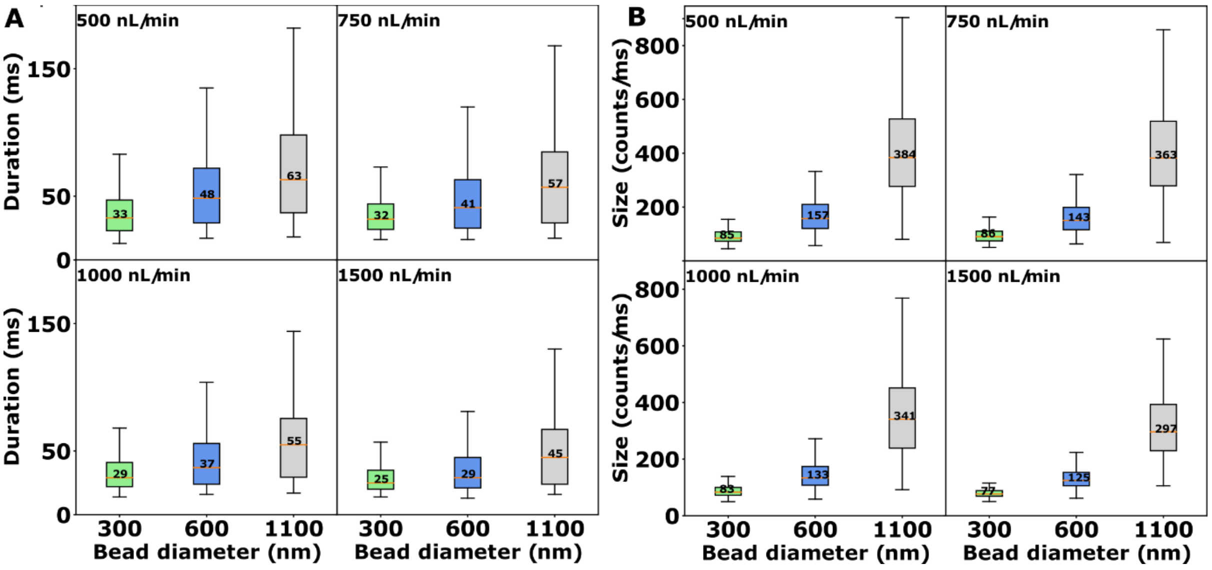


**Fig. S2**. **Sizes and durations of nonspecific signal dips from measurements of unlabeled beads.** The durations (**A**) and sizes (**B**) of the dips in the nonspecific signal from measurements of beads with diameters 300 ,600, and 1,100 nm, traversing the probe volume at flow rates 500, 750, 1000 and 1500 nL/min ($1.39 \times 10^{-6}$, $2.08 \times 10^{-6}$, $2.77 \times 10^{-6}$ and $4.16 \times 10^{-6}$ m/s calculated mean velocities in microfluidic channel, respectively). The showed data is an accumulation from different repeats, see Fig. S5 for details. The orange horizontal line represents the median value.

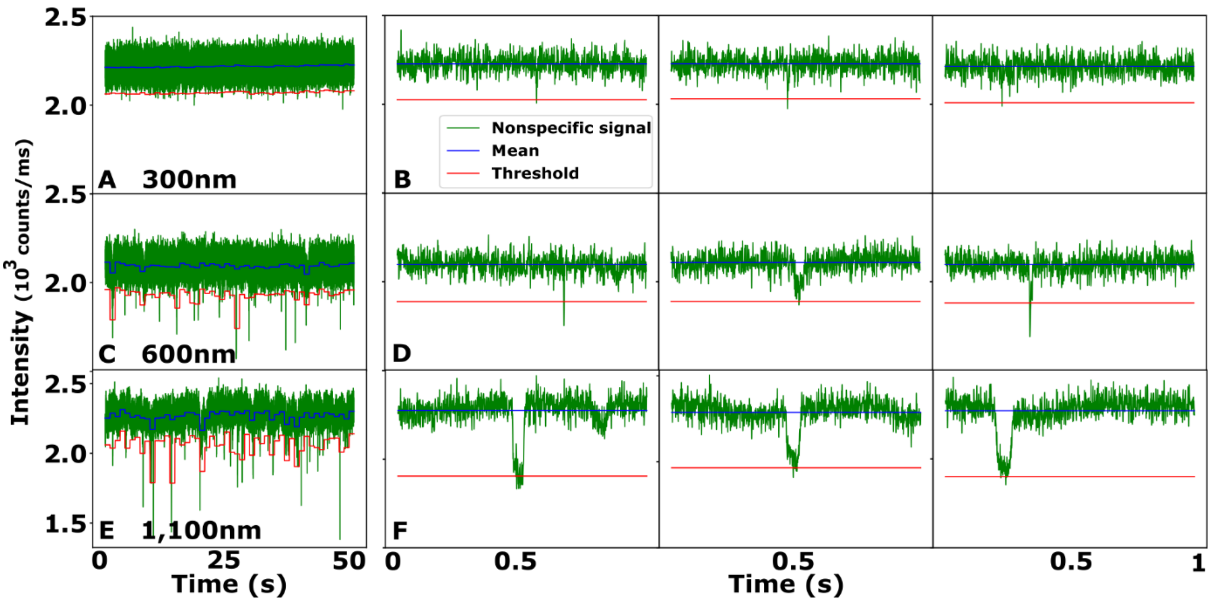


**Fig. S3. Example time trace of measurements of unlabeled beads.** A time window of 50 s from measurement of beads with diameters of **A.** 300 nm, **C.** 600 nm, and **E.** 1,100 nm traversing the probe volume. A time window of 1 s from measurement of beads with diameters of **B.** 300 nm **D.** 600 nm **F.** 1,100 nm traversing the probe volume. Green - nonspecific signal, blue - mean of each 1 s of time window, red - threshold of 4 std below the mean signal of 1 s acquisition for dip determination. All measurements were conducted in the presence of 500 µM Fluorescein with flow rate of 750 nL/min ($2.08 \times 10^{-6}$ m/s calculated mean velocity in microfluidic channel Eq. S1)

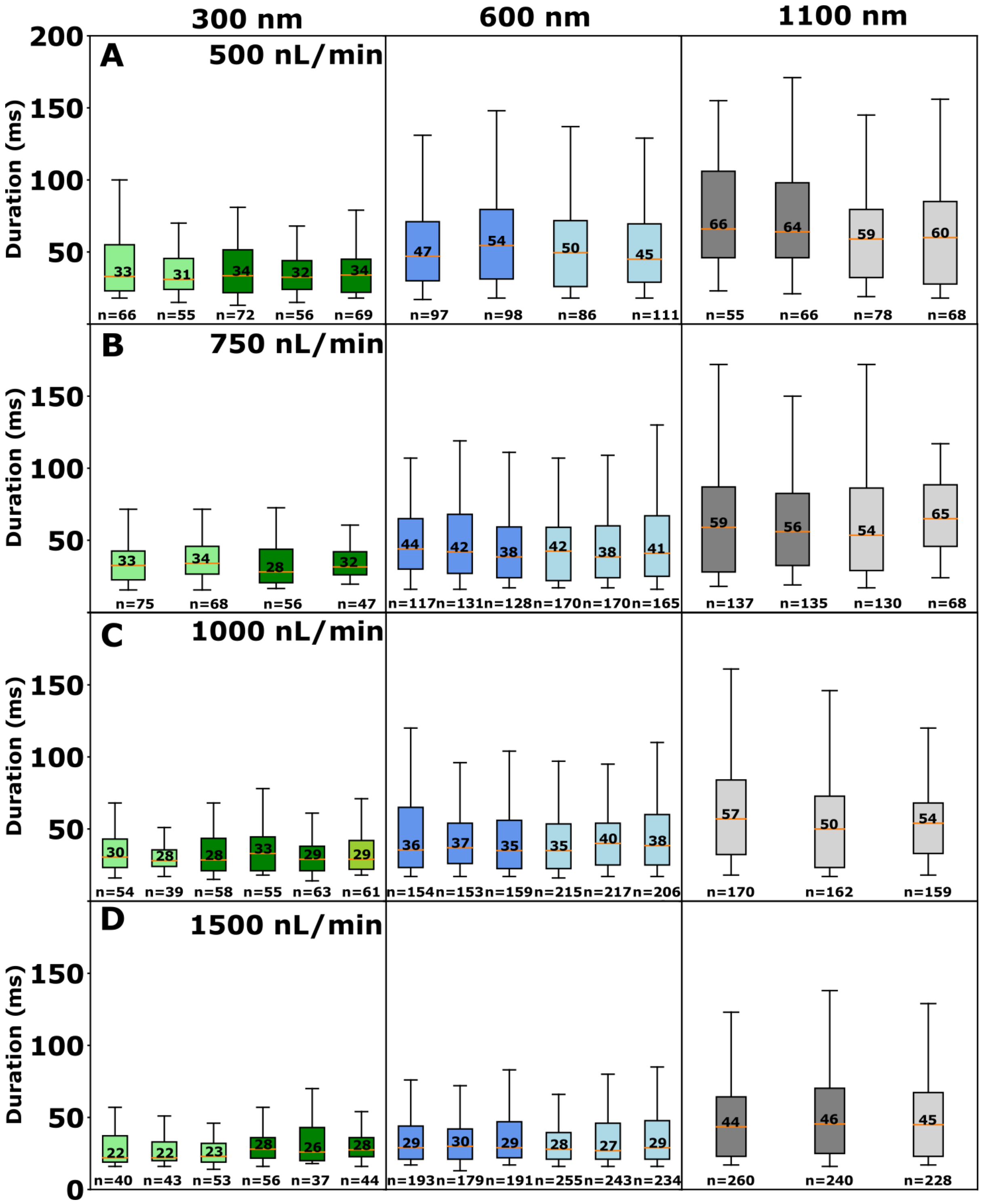


**Fig. S4. Repetitions measurements of unlabeled beads.** Distributions of signal durations of beads with diameters of 300 nm (left column), 600 nm (middle column), and 1,100 nm (right column), in the presence of 500 µM Fluorescein with flow rate of **A.** 500 nL/min ($1.39x10^{-6}$ m/s calculated mean velocity in microfluidic channel) **B.** 750 nL/min ($2.08x10^{-6}$ m/s) **C.** 1000 nL/min ($2.77x10^{-6}$ m/s) **D.** 1500 nL/min ($4.16x10^{-6}$ m/s) from 7 min. acquisitions. Same color represents technical repeats. The orange horizontal line represents the median value. **n=** number of detection events.

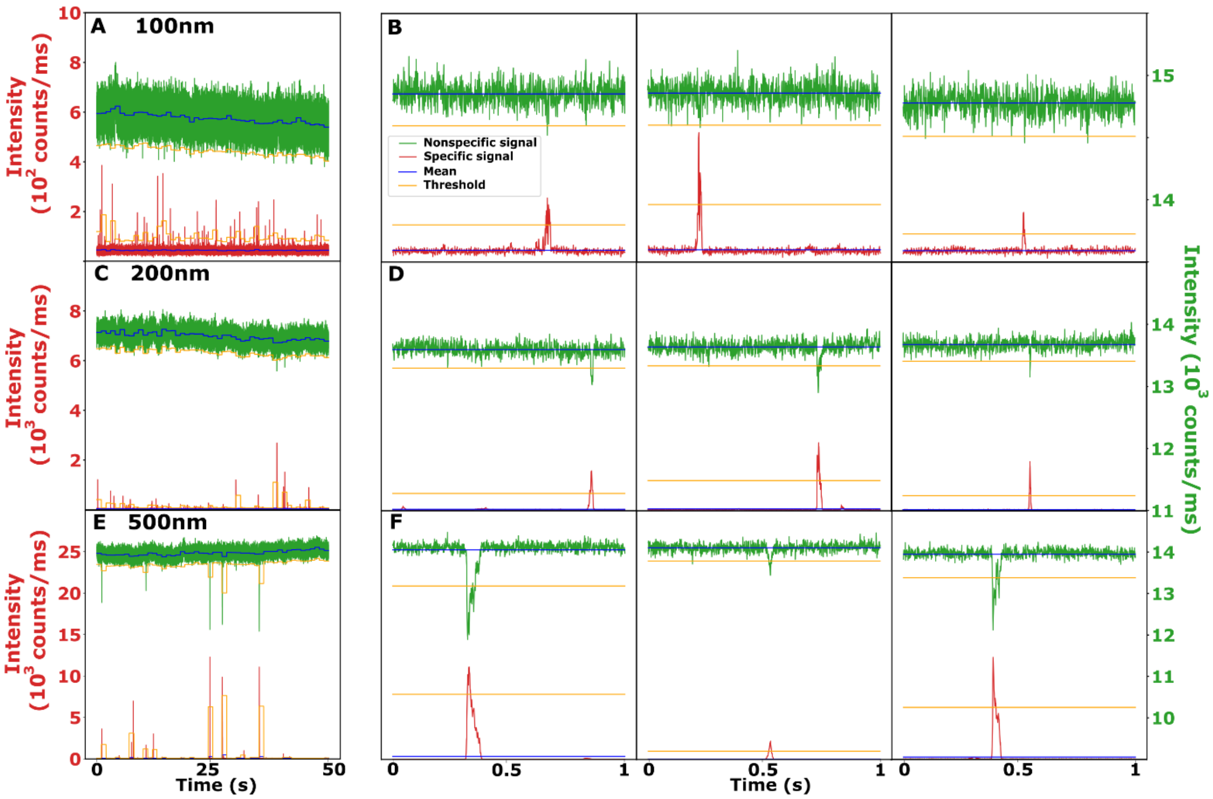


**Fig. S5. Example time traces of labeled beads measurements.** A time window of 50 s from measurement of beads with diameters of **A.** 100 nm, **C.** 200 nm, and **E.** 500 nm traversing the probe volume. A time window of 1 s from measurement of beads with diameters of **B.** 100 nm, **D.** 200 nm, and **F.** 500 nm traversing the probe volume. Green - nonspecific signal, blue - mean of 1 s measurement time, orange - threshold of 3 std below the mean of 1 s measurement time for dip and - threshold of 5 std above the mean of 1 s measurement time for burst. All measurements were conducted in the presence of 500 µM Fluorescein with flow rate of 750 nL/min ($2.08 \times 10^{-6}$ m/s calculated mean velocity in microfluidic channel Eq. S1)

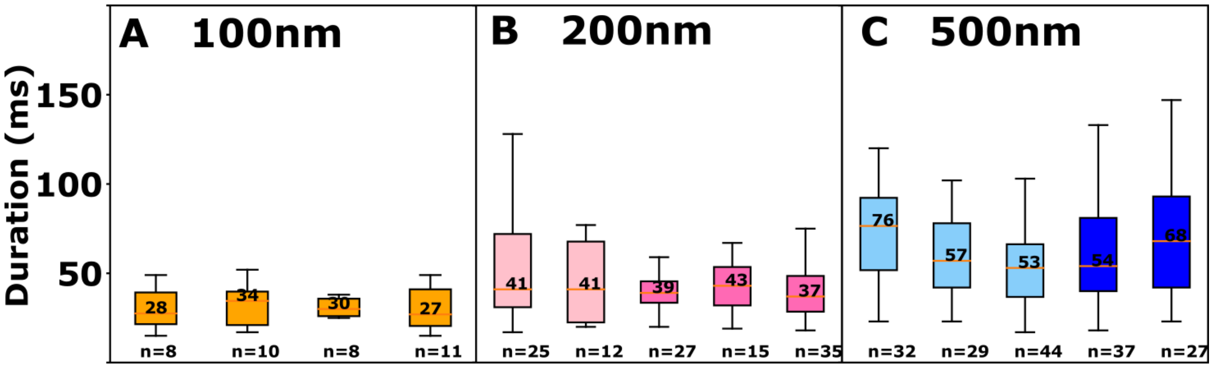


**Fig. S6. Repetitions measurements of labeled beads.** Distributions of signal dip duration of beads with diameters **A.** 100 nm, **B.** 200 nm, and **C.** 500 nm in the presence of 500 µM Fluorescein with flow rate of 750 nL/min ($2.08 \times 10^{-6}$ m/s calculated mean velocity in microfluidic channel, Eq. S1) from 5 min. acquisitions. Same color represents technical repeats. The orange horizontal line represents the median value. **n=** detection events

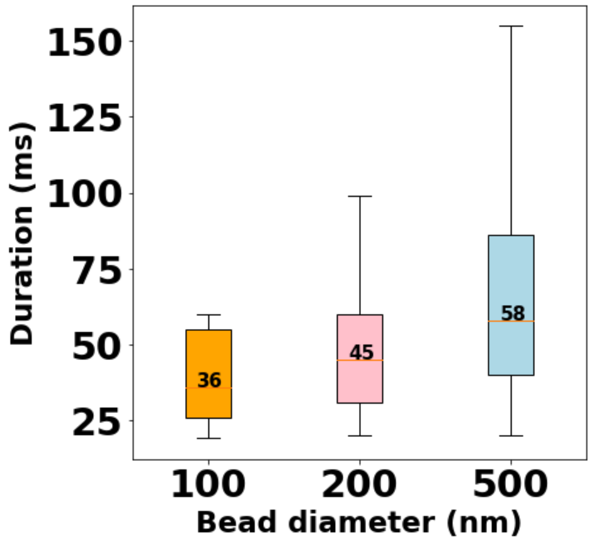


**Fig. S7. Burst durations from coincident burst-dip pairs.** Burst durations of coincident dip-burst pairs with dips (Fig. 3C) from dye-labeled red beads measured in the 3D-printed setup. (accumulation of 37, 114 and 169 detection events from 4, 5 and 5 repetitions of 5 min. acquisitions, respectively). All measurements were conducted in the presence of 500 µM Fluorescein with flow rate of 750 nL/min ($2.08 \times 10^{-6}$ m/s calculated mean velocity in microfluidic channel Eq. S1) The orange horizontal line represents the median value.

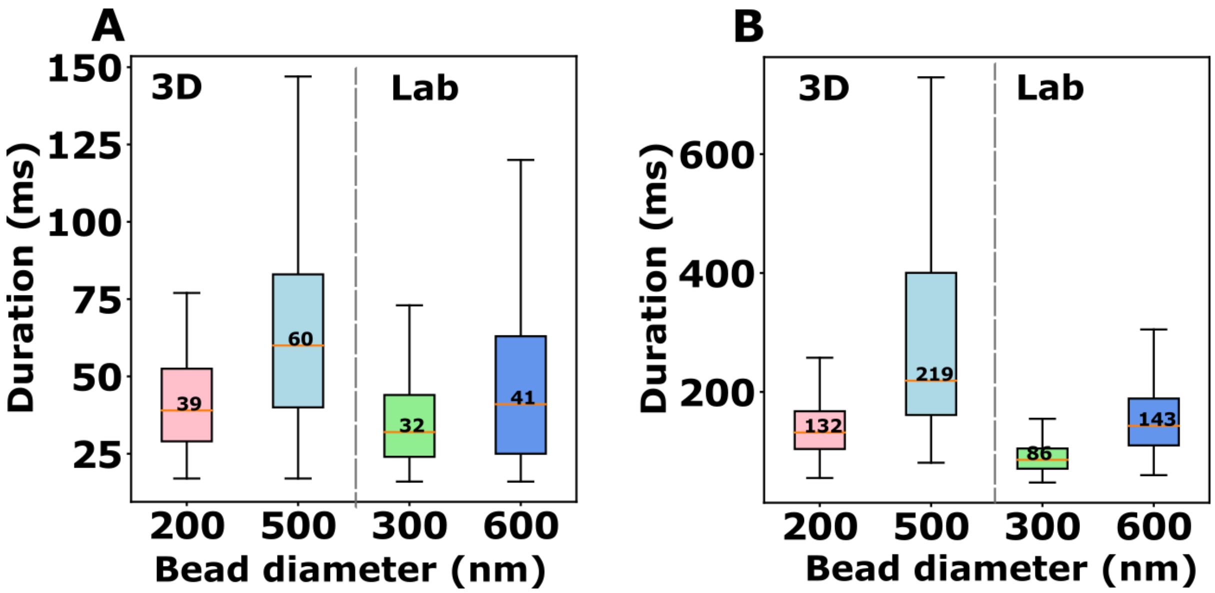


**Fig. S8. Comparison between sizes and durations of dips in the nonspecific signal measured in two experimental setups.** Durations (**A**) and sizes (**B**) of signal dips from measurements of red dye-labeled beads with diameters of 200 nm and 500 nm (5 repetitions of 5 min. acquisitions, see Fig. S6) on the 3D-printed setup, Brick-MIC,(labeled as **3D**) and for unlabeled polyester beads with diameters of 300 nm and 600 nm (4 and 6 repetitions of 7-min. measurements, respectively, see Fig S4B) on the laboratory confocal-based setup (labeled as **Lab**). All measurements were performed at 750 nL/min flow rate ($2.08 \times 10^{-6}$ m/s calculated mean velocity in microfluidic channel) in the presence of 500 µM Fluorescein. The orange horizontal line represents the median value.

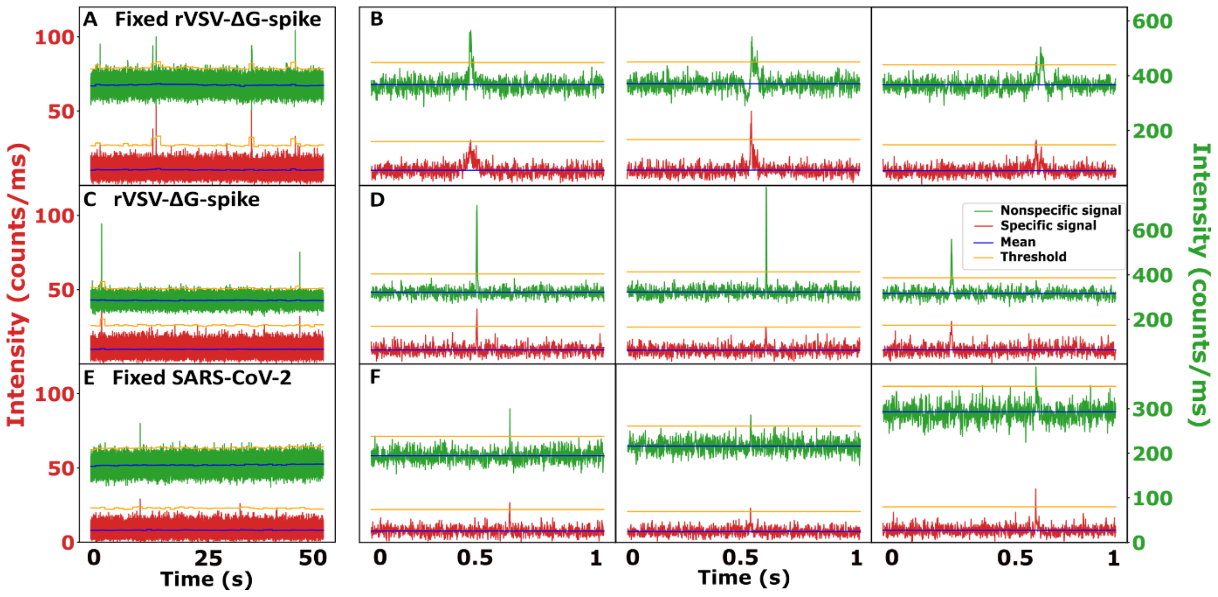


**Fig. S9. Examples of traces from virus measurements.** A time window of 50 s from measurement of **A.** fixed rVSV-ΔG-spike **C.** non-fixed rVSV-ΔG-spike, and **E.** fixed SARS-CoV-2 traversing the probe volume. A time window of 1 s from measurement of **B.** fixed rVSV-ΔG-spike, **D.** non-fixed rVSV-ΔG-spike, and **F.** fixed SARS-CoV-2 traversing the probe volume. Green - nonspecific signal, blue - mean of 1 s measurement time, orange - threshold of 3 std above the mean of 1 s measurement time for burst in the nonspecific signal and - threshold of 5 std above the mean of 1 s measurement time for burst in the specific signal. All measurements were performed at 750 nL/min flow rate ($2.08x10^{-6}$ m/s calculated mean velocity in microfluidic channel, Eq. S1) in the presence of eq. 10 µM fluorescein-labeled BSA.

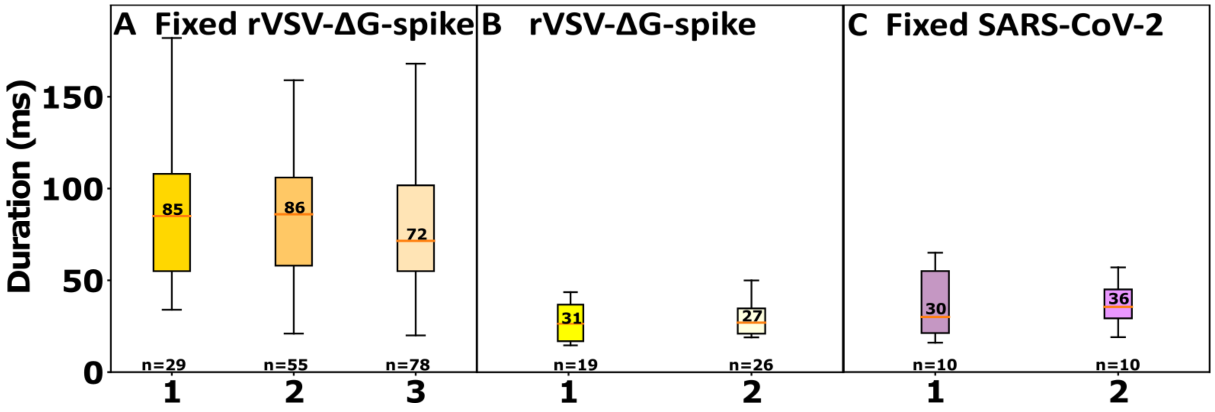


**Fig. S10. Biological repeats of virus measurements.** Biological repeat for **A.** fixed rVSV-ΔG-spike:1-six technical repeats, 2- five technical repeats, 3- nine technical repeats. **B.** live rVSV-ΔG-spike:1-three technical repeats, 2- three technical repeats. **C.** fixed SARS-CoV-2: 1-five technical repeats, 2- ten technical repeats. All measurements (each technical repeat) were conducted for 5 min. acquisitions at 750 nL/min flow rate ($2.08x10^{-6}$ m/s calculated mean velocity in microfluidic channel) with eq. 10 µM fluorescein-labeled BSA, **n**=accumulation of detection from all the measurement repeats. The orange horizontal line represents the median value.

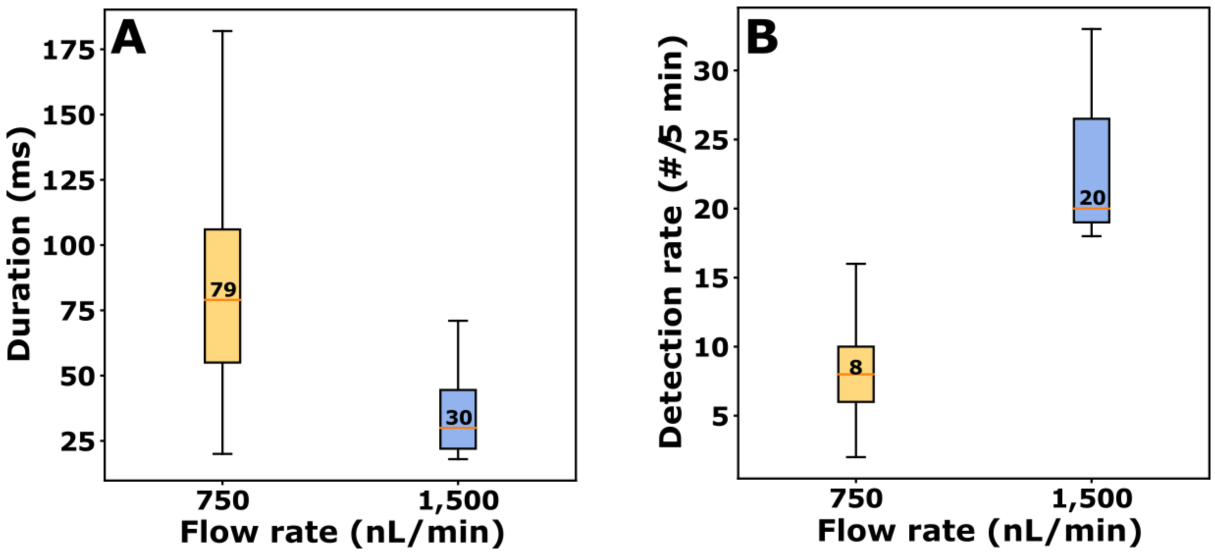


**Fig. S11. Measurement of fixed and neutralized rVSV-ΔG-spike at different flow rates. A.** Burst durations from a 5 min. acquisition in different flow rates 750 and 1500 nL/min ($2.08x10^{-6}$ and $4.16x10^{-6}$ m/s calculated mean velocity in microfluidic channel, Eq. S1), accumulation of 157 and 71 detection events from 18 and 3 repetitions, respectively. **B.** 8.7±1.0 (SEM) and 23.6±3.8 (SEM) detection events from 5 min. acquisitions. All measurements were performed in the presence of eq. 10 μM Fluorescein-labeled BSA at flow rate of 750 or 1500 nL/min ($2.08x10^{-6}$ or $4.16x10^{-6}$ m/s calculated mean velocity in microfluidic channel, Eq. S1)

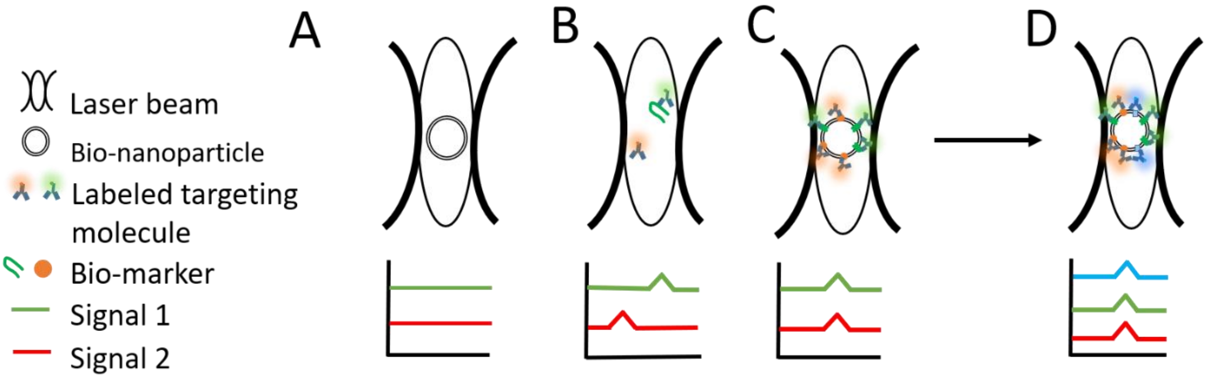


**Fig. S12. Alternative detection layouts. A**. Unlabeled bio-nanoparticle does not lead to a change in the nonspecific detection channel. **B** Dye-labeled antibodies lead to uncorrelated bursts only in the specific signal detection channel. **C** Doubly labeled bio-nanoparticle simultaneously lead to a double burst in the specific signal detection channels, with burst duration proportional to the bio-nanoparticle volume. **D** Option C can be extended for the use of three differently-labeled different antibodies targeting three different targets on top of the same bio-nanoparticle.

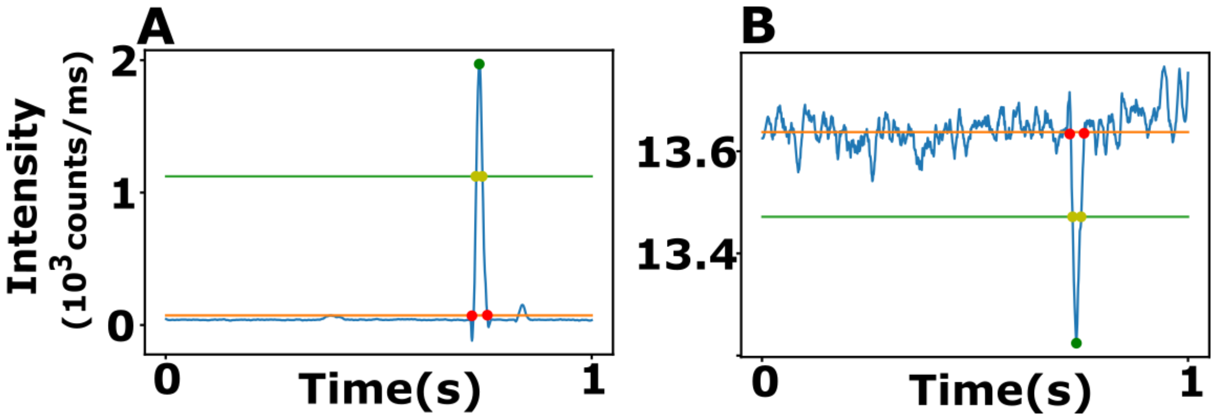


**Fig. S13. Example of calculation of the signal change characteristics A.** burst **B.** dip. Blue line - smoothed data of 1,000 data points frame (equal to 1 s). Green line – threshold line equal of the mean of the data frame ± number of std (see Table S5). Light green dot - intersecting time point between green and blue lines. Dark green dot – extremum point between two intersecting time points (light green dot) between green and blue lines. Orange line - equal to the mean of the data frame. Red dots - data points that are equal to the intersecting time point between orange and blue lines, defined as the initial and final time points of the burst/dip.

### Supplementary Tables

**Table S1. P-values of two-sample t-tests**

| **Figure1, Panel1**<br><br>**or**<br><br>**Figure1, Panel1 vs.**<br>**Figure2, Panel2** | **Sample #1** | **Sample #2** | **P-value** | **Significance** |
|---|---|---|---|---|
| 2B | 300 nm[1] dip duration | 600 nm dip duration | $1.3x10^{-10}$ | ***[5] |
| | 600 nm dip duration | 1,100 nm dip duration | $1.8x10^{-20}$ | *** |
| 2C | 300 nm dip size | 600 nm dip size | $6.8x10^{-37}$ | *** |
| | 600 nm dip size | 1,100 nm dip size | $1.9x10^{-186}$ | *** |
| 3C, 3D | 100 nm burst duration | 100 nm dip duration | 0.22 | n.s[2] |
| | 200 nm burst duration | 200 nm dip duration | 0.08 | n.s |
| | 500 nm burst duration | 500 nm dip duration | 0.97 | n.s |
| 3C | 100 nm dip duration | 200 nm dip duration | 0.02 | *[3] |
| | 200 nm dip duration | 500 nm dip duration | $2.3x10^{-8}$ | *** |
| 3D | 100 nm burst duration | 200 nm burst duration | $6.6x10^{-22}$ | *** |
| | 200 nm burst duration | 500 nm burst duration | $3.7x10^{-25}$ | *** |
| 4B | Fixed rVSV-ΔG-spike burst duration | 500 nm burst duration | $7.4x10^{-9}$ | *** |
| 4D | rVSV-ΔG-spike burst duration | 100 nm burst duration | 0.06 | n.s |
| 4F | SARS-CoV-2 burst duration | 100 nm burst duration | 0.65 | n.s |
| | SARS-CoV-2 burst duration | 200 nm burst duration | 0.04 | * |
| S1C | 300 nm burst duration flow | 300 nm burst duration no flow | $7.0x10^{-19}$ | *** |
| S7 | 100 nm dip duration | 200 nm dip duration | 0.53 | n.s |
| | 200 nm dip duration | 500 nm dip duration | $2.06x10^{-6}$ | *** |
| S7, 3C | 100 nm burst duration | 100 nm dip duration | 0.07 | n.s |
| | 200 nm burst duration | 200 nm dip duration | 0.25 | n.s |
| | 500 nm burst duration | 500 nm dip duration | 0.98 | n.s |
| S7, 3D | 100 nm burst duration | 100 nm burst duration | 0.22 | n.s |
| | 200 nm burst duration | 200 nm burst duration | 0.67 | n.s |
| | 500 nm burst duration | 500 nm burst duration | 0.95 | n.s |

| | | | | |
|---|---|---|---|---|
| S10A | Repeat 1 | Repeat 2 | 0.92 | n.s |
| | Repeat 1 | Repeat 3 | 0.43 | n.s |
| | Repeat 2 | Repeat 3 | 0.4 | n.s |
| S10B | Repeat 1 | Repeat 2 | 0.14 | n.s |
| S10C | Repeat 1 | Repeat 2 | 0.89 | n.s |

[1] – Bead diameter
[2] – n.s → $p>0.05$
[3] – * → $0.01<p<0.05$
[4] – ** → $0.001<p<0.01$
[5] – *** → $p<0.001$

**Table S2. Control for specific detection of dye-labeled beads.** For control, the function *'durationsizecross'* was used for detecting coincident detection events as changes in the specific and nonspecific signals (dips/bursts).

| Sample | Rate of coincident detection events (per min.) |
|---|---|
| Background[1] | 0 |
| 600 nm diameter unlabeled[2] | 0 |

[1] - Background, is the measurement of solely free dyes, without any nanoparticles (m=4)
[2] – Beads of 600 nm diameter (m=4).
m= No. of technical repeats
All measurements were conducted in the presence of 500 µM fluorescein with microfluidic laminar flow at 750 nL/min flow rate ($2.08 \times 10^{-6}$ m/s calculated mean velocity in microfluidic channel).

**Table S3. Quantification of false positive coincident detection events due to leakage of fluorescence from the nonspecific signal bursts to the specific signal detection channel at different free dye concentrations.** As the concentration of the free dye is reduced, the number of false positive events are reduced.

| Free dye concentration (µM) | Average number of false positive events per 5 min. acquisition (± SEM) |
|---|---|
| 100 | 9.3±1.4 |
| 50 | 2.0±0.9 |
| **10** | **0.1±0.1**[1] |

All measurements were conducted in the present of free dye and fixed rVSV-ΔG-spike at 750 nL/min flow rate ($2.08 \times 10^{-6}$ m/s calculated mean velocity in microfluidic channel) for 5 min. acquisition at concentration of 10 µM n=10, concentration 50 µM n=4, concentration 100 µM n=3. n= No. of repeats

[1] – bolded to highlight the fact that we used this condition in our measurements.

| **Table S4. Control for specific detection of fixed rVSV-ΔG-spike**. As a control, we analyzed data from different measured conditions for the quantifying coincident detection events between the specific and non-specific signal detection channels. All acquisitions were conducted in the presence of 10 µM fluorescein-labeled BSA in 750 nL/min flow rate (2.08x$10^{-6}$ m/s calculated mean velocity in microfluidic channel) for 5 min. acquisition (Table S2, bolded). | | | | | | | |
|---|---|---|---|---|---|---|---|
| **Mixture constituents** | | | | | | **Average coincident events per 5 min. acquisition (± SEM)** | **Number of repeats** |
| **Condition ID** | **Free dye** | **Human serum albumin (HSA)** | **virus** | **Anti-spike labeled Ab** | **Nonspecific labeled Ab** | | |
| A | X | | | | | 0.14±0.01 | 14 |
| B | X | X | | | | 0.17±0.01 | 17 |
| C | X | X | X | | | 0.10±0.10 | 10 |
| D | X | X | | X | | 0.14±0.08 | 21 |
| E | X | X | X | | X | 0.10±0.10 | 10 |
| **F**[1] | **X** | **X** | **X** | **X** | | **8.72±1.02** | **18** |

A- Background, measurement of only free dye without virus
B- ~10 µM Free BSA-fluorescein and HSA
C- Fixed rVSV-ΔG-spike in the present of ~10 µM free BSA-fluorescein and HSA
D- ~10 µM Free BSA-fluorescein and HSA and non-specific labeled
E- Fixed rVSV-ΔG-spike in the present of ~10 µM free BSA-fluorescein, HSA and non-specific labeled
F-Fixed rVSV-ΔG-spike in the present of ~10 µM free BSA-fluorescein, HSA and labeled antibodies

[1] – bolded to highlight the fact that we used this condition in our measurements.

| **Table S5. Thresholds for finding a dip/burst** | | |
|---|---|---|
| Sample | Nonspecific signal | Specific signal |
| Unlabeled beads | 4 std below the mean | - |
| Labeled beads | 3 std below the mean | 5 std above the mean |
| Virus | 3 std above the mean | 5 std above the mean |
